# Charge Screening by Extracellular Ca²⁺ Shields Endothelial Cells from Histone Injury and Permits Membrane Repair

**DOI:** 10.1101/2025.07.13.664578

**Authors:** Jade H. Cleary, Adrian M. Sackheim, Rushi Sonawane, Taylor Seitz, Bryan Walters, Kara J. Wentzel, Abigail J. Howard, Lydia Sheeser, Rachel Barba, Sophia H. Piffard, J. Cory Benson, Mohamed Trebak, Susan M. Shea, Matthew D. Neal, Nikita Mohapatra, Aijun Wang, Lucy Z. Kornblith, Mitchell J. Cohen, Michael B. Yaffe, Devdoot Majumdar, Grant Hennig, Mark T. Nelson, Kalev Freeman

## Abstract

**Background:** Extracellular histones released after trauma and in sepsis injure the vascular endothelium and contribute to coagulopathy and organ failure. Hypocalcemia frequently accompanies resuscitation with citrated blood products and predicts poor outcome, an association usually attributed to coagulation-factor dysfunction. Whether extracellular Ca²⁺ directly governs endothelial vulnerability to circulating cytotoxic histones, and whether the resulting membrane injury is reparable, are unknown.

**Methods:** We combined live-cell spinning-disk confocal imaging of plasma membrane dynamics (FM1-43), Ca²⁺ imaging, flow cytometry, electric cell-substrate impedance sensing, and protein-lipid binding assays in cultured human endothelial cells, ORAI1/2/3 triple-knockout HEK-293 cells and surgically opened *en face* murine mesenteric arteries. Membrane repair was assessed with a sequential two-dye live-cell viability assay.

**Results:** Histones evoked rapid, irregular endothelial Ca²⁺ elevations that were unaffected by deletion of all three ORAI homologs and were blocked by Gd³⁺ only at concentrations two orders of magnitude above those required to inhibit store-operated Ca²⁺ entry, excluding this pathway. Histones instead bound anionic membrane phospholipids, deforming the plasma membrane: lamellipodial ruffling, retraction, blebbing, extracellular vesicle release, phosphatidylserine externalization, and permeabilization in a heterogeneous subset of cells. Removing extracellular Ca²⁺ did not prevent these changes but amplified them, increasing dye-permeable cells from 34±7% to 60±11% and converting the initial rise in monolayer electrical resistance into immediate barrier failure. Conversely, elevated Ca²⁺ (12 mM) or Gd³⁺ (10 µM) prevented membrane deformation, while high Ca²⁺ markedly reduced and Gd³⁺ (30 µM) abolished histone binding to those phospholipids. A subset of permeabilized cells subsequently restored membrane integrity, whereas Ca²⁺-overloaded cells never recovered.

**Conclusions:** Extracellular Ca²⁺ limits histone-induced endothelial injury through a mechanism consistent with electrostatic screening of histone–membrane interactions, rather than through its contribution to intracellular Ca²⁺ overload. Histone-induced membrane permeabilization is reparable in a subpopulation of endothelial cells, identifying membrane repair as a candidate therapeutic target.

## Introduction

The more than 60 trillion interconnected endothelial cells (ECs) that line the vasculature regulate blood flow, coagulation, and barrier function.^1^ In critical illness such as trauma and sepsis, shock-induced endotheliopathy contributes to progressive organ failure and death.^2^ Release of cytotoxic histone proteins into the circulation, from injured cells and from neutrophil extracellular traps, drives this acute endothelial injury.^3–9^ The mechanisms by which histones injure the endothelium remain incompletely defined, which impedes treatment design.

Two features of this problem have direct clinical consequences. The first is hypocalcemia, which occurs commonly during resuscitation of critically injured patients through infusion of citrated blood products and reduced hepatic citrate clearance.^10–13^ Low ionized Ca²⁺ predicts mortality and the need for multiple transfusions, but thresholds, timing, and extent of repletion remain contested, in part because hypocalcemia has been framed almost exclusively as a coagulation-factor problem. Whether extracellular Ca²⁺ acts directly on the endothelium to set its vulnerability to circulating cytotoxic mediators has not been tested. The second is the fate of injured endothelium. Clinical trials showing that prehospital transfusion of donor plasma reduces biomarkers of endothelial injury and improves survival imply that the injured endothelium retains capacity to stabilize or recover.^14^ Yet whether an endothelial cell that has lost its plasma membrane integrity can repair itself is unknown.

We previously showed that histones initiate large endothelial Ca²⁺ oscillations that are pro-inflammatory, pro-thrombotic, and degrade barrier function, but the route of Ca²⁺ entry was unknown.^7,15,16^ Store-operated Ca²⁺ entry (SOCE), mediated by STIM1-gated ORAI channels, is the dominant Ca²⁺ influx pathway in non-excitable cells including ECs, and is activated by stimuli such as VEGF and thrombin.^17–21^ We therefore first hypothesized that SOCE contributes to histone-induced Ca²⁺ entry. We also considered an alternative: that cationic histones interact electrostatically with anionic phospholipids on the endothelial surface, deforming the membrane and permitting Ca²⁺ entry directly. This is plausible because histones bind phospholipids and disrupt lipid bilayers.^22–24^ This alternative model has a counterintuitive and clinically important corollary: because extracellular Ca²⁺ screens membrane surface charge,^25,26^ *lowering* Ca²⁺ should increase the negative charge density near the endothelial surface and thereby worsen, not lessen, histone toxicity.

We studied ECs in culture and in surgically opened vascular preparations using live-cell video microscopy to address five questions: (1) Are histone-induced Ca²⁺ oscillations SOCE-dependent? (2) Is Ca²⁺ influx necessary for histone-induced membrane injury? (3) Do extracellular Ca²⁺ levels modulate the membrane effects of histones? (4) Do histone-induced membrane changes depend on the negative charge density near the cell surface? (5) Can endothelial cells that have been permeabilized by histones repair their plasma membranes?

## Methods

A detailed description of all experimental procedures is provided in the Supplemental Material. Human blood was collected from healthy adult volunteers who gave written informed consent, under a protocol approved by the University of Vermont Institutional Review Board; all animal studies were approved by the University of Vermont Institutional Animal Care and Use Committee and conducted in accordance with the NIH Guide for the Care and Use of Laboratory Animals. Experiments in mice used male animals only, and sex-based differences were therefore not assessed. Data were tested for normality using the Kolmogorov-Smirnov test, and parametric or non-parametric comparisons applied accordingly, with significance at P<0.05.

## Results

### Store-operated Ca²⁺ entry does not contribute to histone-induced endothelial Ca²⁺ events

To test whether SOCE contributes to histone-induced Ca²⁺ elevations, we first used Gd³⁺, a trivalent lanthanide that inhibits SOCE at low concentrations (<10 µM).^23^ Freshly harvested murine mesenteric arteries, surgically opened to expose the endothelium *en face*, were loaded with a fluorescent Ca²⁺ indicator and exposed to clinically relevant concentrations of histones (50 µg/mL) in 1.2 mM Ca²⁺ with or without Gd³⁺ (10 µM) (Figure 1A–C; Supplemental Videos 1, 2). Rapid (<5 min), irregular increases in endothelial Ca²⁺ fluorescence were observed, as expected. These oscillations were recapitulated in HEK-293 cells loaded with Fura-2: Ca²⁺ signals appeared at histone concentrations of 25 µg/mL, with high-amplitude oscillations at 50 µg/mL (Figure 1D, E). As we previously demonstrated in vascular preparations,^7^ extracellular Ca²⁺ was required; histones (50 µg/mL) applied in 0 mM Ca²⁺ produced no Ca²⁺ activity, and restoring extracellular Ca²⁺ to 2 mM after 5 min produced an immediate increase in activity (Supplemental Figure 1C).

**Figure 1.**
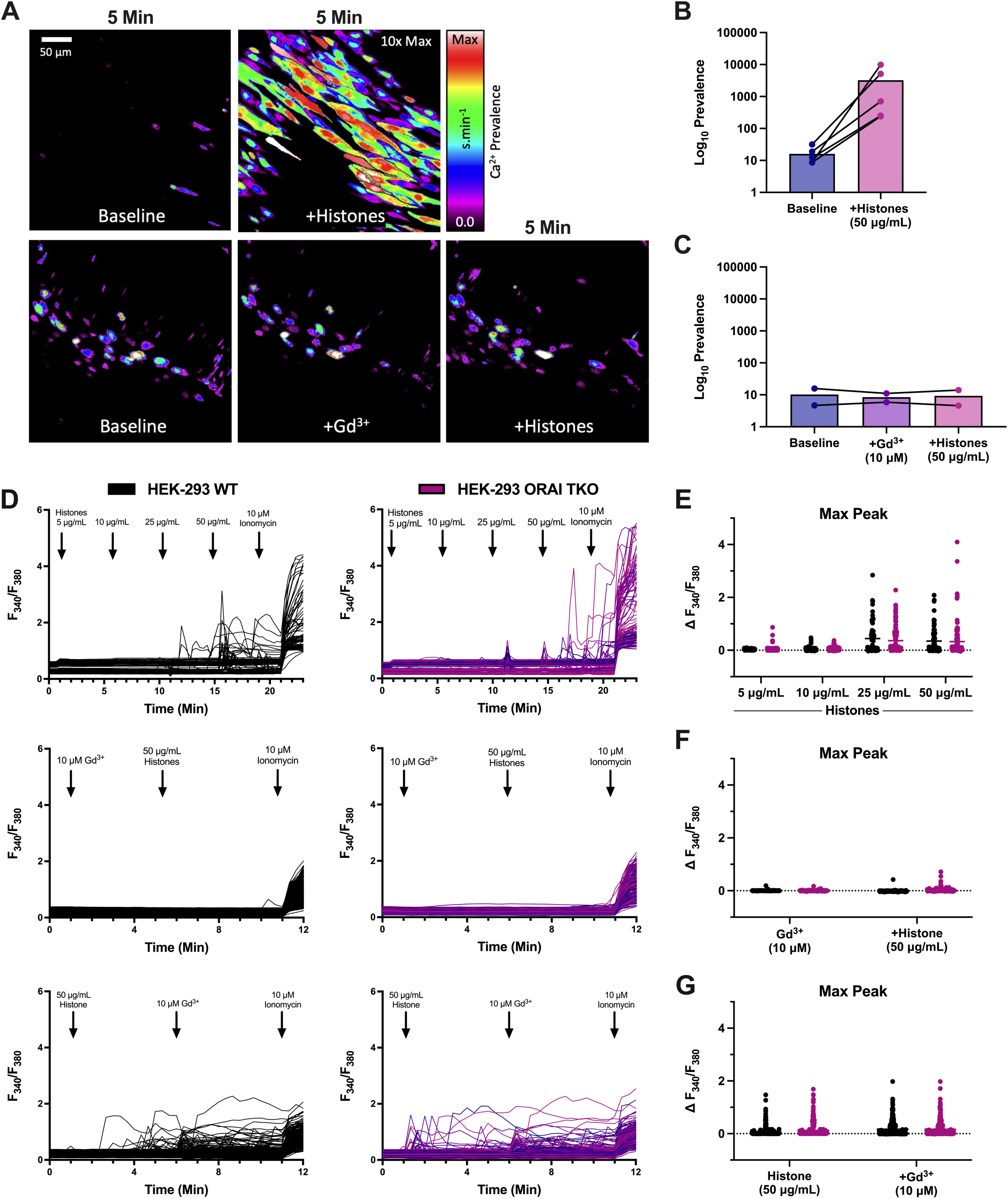
Histone-induced endothelial Ca²⁺ influx is independent of ORAI channels and blocked by Gd³⁺ only at charge-screening concentrations. (**A**) *En face* murine mesenteric arteries stimulated with histones (50 µg/mL) with or without Gd³⁺ (10 µM) pre-treatment. Cumulative Ca²⁺ prevalence images at baseline and after 5 min of histones (top), or after Gd³⁺ pre-treatment followed by histones (bottom). (**B**) Ca²⁺ activity at baseline versus histone stimulation. (**C**) Ca²⁺ activity at baseline, after Gd³⁺ pre-treatment, and during subsequent histone stimulation. (**D**) Fura-2 traces in wild-type (left) and ORAI triple-knockout (right) HEK-293 cells during increasing histone concentrations (top), Gd³⁺ pre-treatment followed by histones (middle), and Gd³⁺ post-treatment (bottom). (**E–G**) Maximum peak intensity per cell for histone titration (**E**), Gd³⁺ pre-treatment (**F**), and Gd³⁺ post-treatment (**G**). Acquisition, quantification, and statistics in Supplemental Methods.

Pre-treatment with Gd³⁺ (10 µM) completely blocked histone-induced Ca²⁺ signals (Figure 1D, F), whereas post-treatment within 5 min of histone exposure did not (Figure 1D, G). A lower Gd³⁺ concentration (5 µM), which should inhibit SOCE-dependent activity,^19^ failed to block histone-induced Ca²⁺ increases in either configuration (Supplemental Figure 1A, B). Because the IC₅₀ of Gd³⁺ for SOCE is below 100 nM,^27^ a requirement for ≥10 µM argues against a SOCE mechanism.

To rule out SOCE definitively, we used a genetic approach.^28^ ORAI channels mediate SOCE by binding STIM1 to open the Ca²⁺ release-activated Ca²⁺ channel,^29^ and deleting all three ORAI homologs abolishes functional SOCE.^28,30,31^ There was no difference in histone-induced Ca²⁺ activity between wild-type and ORAI triple-knockout (TKO) HEK-293 cells, and Gd³⁺ still blocked responses in TKO cells (Figure 1D–G). ORAI channels therefore do not contribute to histone-induced Ca²⁺ influx.

### Histones bind and deform endothelial plasma membranes

Our observations that histone-induced Ca²⁺ entry was ORAI-independent, and that Gd³⁺ blocked it only above 5 µM, suggested Gd³⁺ acts through a different mechanism. Multivalent cations such as Gd³⁺ neutralize negative surface charge and alter the local electric field near the membrane.^25,26^ If positively charged histones directly disrupt lipid bilayers,^22–24^ then Gd³⁺ at sufficient concentration could screen the cell surface from cationic histones, preventing the membrane damage that permits Ca²⁺ entry.

Scanning electron microscopy of *en face* murine mesenteric arteries after histones (50 µg/mL, 30 min) showed marked changes from the flat, smooth, junction-intact untreated surface: cells had retracted, cell-to-cell junctions had broken down, retraction fibers were exposed, and numerous blebs and extracellular vesicles were present (Figure 2A).

**Figure 2.**
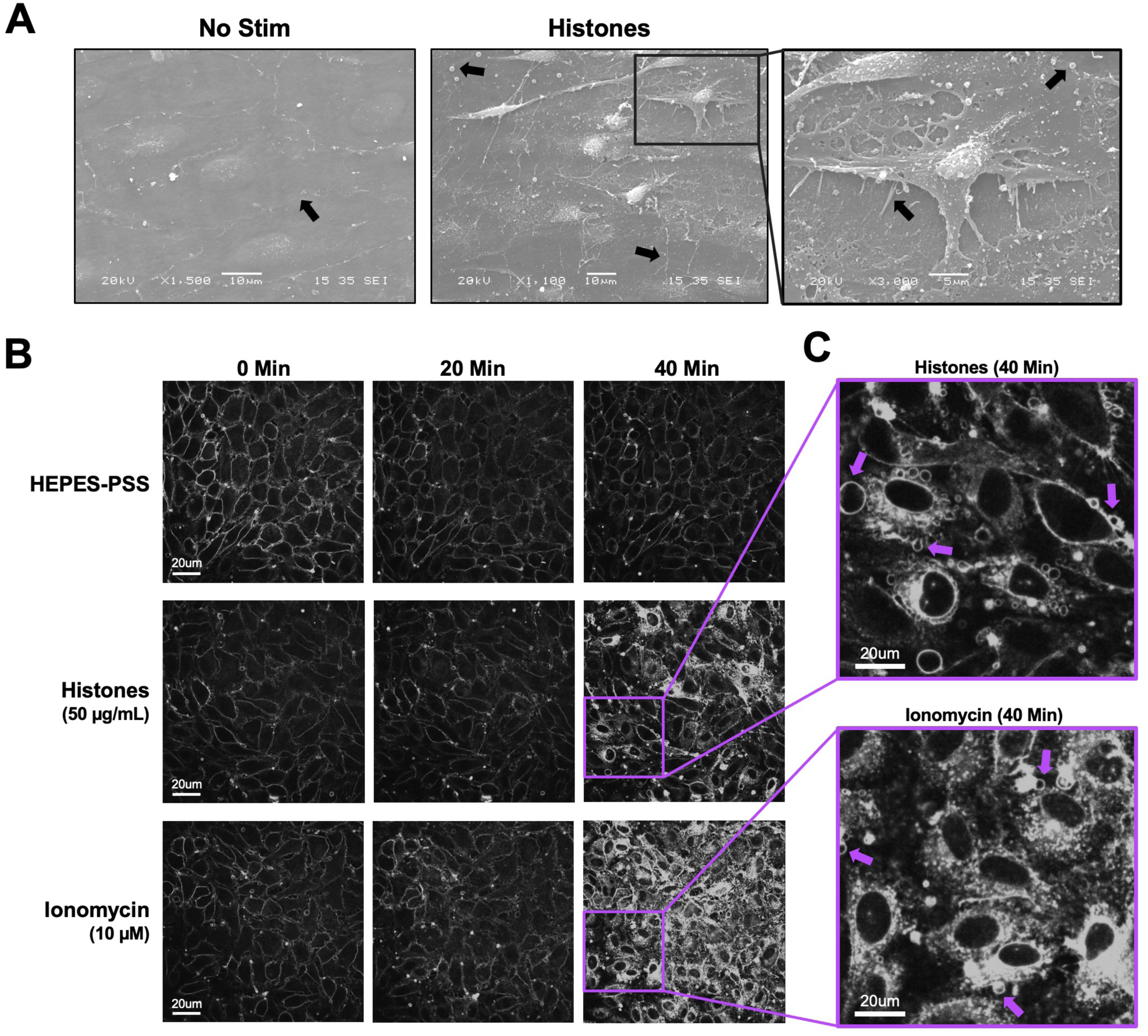
Histones deform endothelial plasma membranes and permeabilize a subset of cells. (**A**) Scanning electron microscopy of native endothelium exposed *en face* in murine mesenteric arteries, untreated (left) and after histones (50 µg/mL, 30 min; right). Arrows indicate retraction fibers, blebs, and extracellular vesicles. (**B**) Representative FM1-43-stained cultured ECs during exposure to buffered saline, histones (50 µg/mL), or ionomycin (10 µM); same field of view at 0, 20, and 40 min. (**C**) Magnification of histone- and ionomycin-stimulated cells at 40 min; arrows indicate blebs.

To visualize these events in live cells, we labeled cultured EC plasma membranes with the styryl pyridinium dye FM1-43, which fluoresces on insertion into lipid bilayers and, being normally cell-impermeant, floods the cytoplasm and stains internal membranes when the membrane is disrupted or aqueous pores open.^32^ We compared histones with ionomycin at a concentration sufficient to produce sustained Ca²⁺ influx.^33^ Both histones and ionomycin permeabilized membranes over 40 min, but differently. Ionomycin produced synchronized cytoplasmic dye filling between 20 and 30 min, with 92±12% of cells showing a ≥2-fold fluorescence increase at 40 min (Figure 2B, C). Histones produced dye internalization in only a subset of ECs (34±7%) at 1.2 mM Ca²⁺; in the remaining approximately two-thirds, dye entry was absent or confined to a subcellular region. Timing was also non-uniform, with few cells responding before 20 min and most internalization occurring between 20 and 40 min, far later than the 2–5 min required for histones to produce Ca²⁺ surges (Figure 1).

Video recordings revealed cell shape and membrane deformation^34^ and extracellular vesicle release in response to both ionomycin and histones, but the patterns of response were qualitatively distinct (Figure 3; Supplemental Videos 3–11). Ionomycin exposure caused filopodial extension and blebbing (Figure 3A). Histones produced cell rounding with exposed retraction fibers and ruffling sheets of plasma membrane resembling lamellipodia at cell edges,^35^ particularly at surfaces adjoining neighboring cells (Figure 3B), together with an end-stage pattern of blebbing and cessation of intracellular trafficking motion in some cells. Vesicle concentrations in the supernatant were higher after 60 min of either agonist than after buffer treatment (buffer, 2.6–4.0 × 10⁷; histones, 4.2–5.2 × 10⁹; ionomycin, 8.6–11 × 10⁹ particles/mL; two independent biological replicates, particles <500 nm). Responses to ionomycin were near-synchronous and symmetrical, with punctate, freely moving particles <5 µm appearing within 10 min, consistent with Ca²⁺-dependent exocytosis of preformed vesicles such as Weibel-Palade bodies.^36,37^ In contrast, histone-induced dye uptake frequently originated at subcellular “hot spots” which coincided with the landing spots for vesicles. We observed membrane activation spreading from these hot spots to the remaining cytoplasm while the nucleus remained unstained, a pattern consistent with endocytic trafficking of labeled lipids. In several recordings an extracellular vesicle approached and contacted the cell surface at the point where a hot spot subsequently appeared, suggesting that these origins mark sites of vesicle attachment rather than arising at random (Figure 8). Membrane remodeling was consistently organized around these endocytic foci rather than distributed evenly across the cell surface.

**Figure 3.**
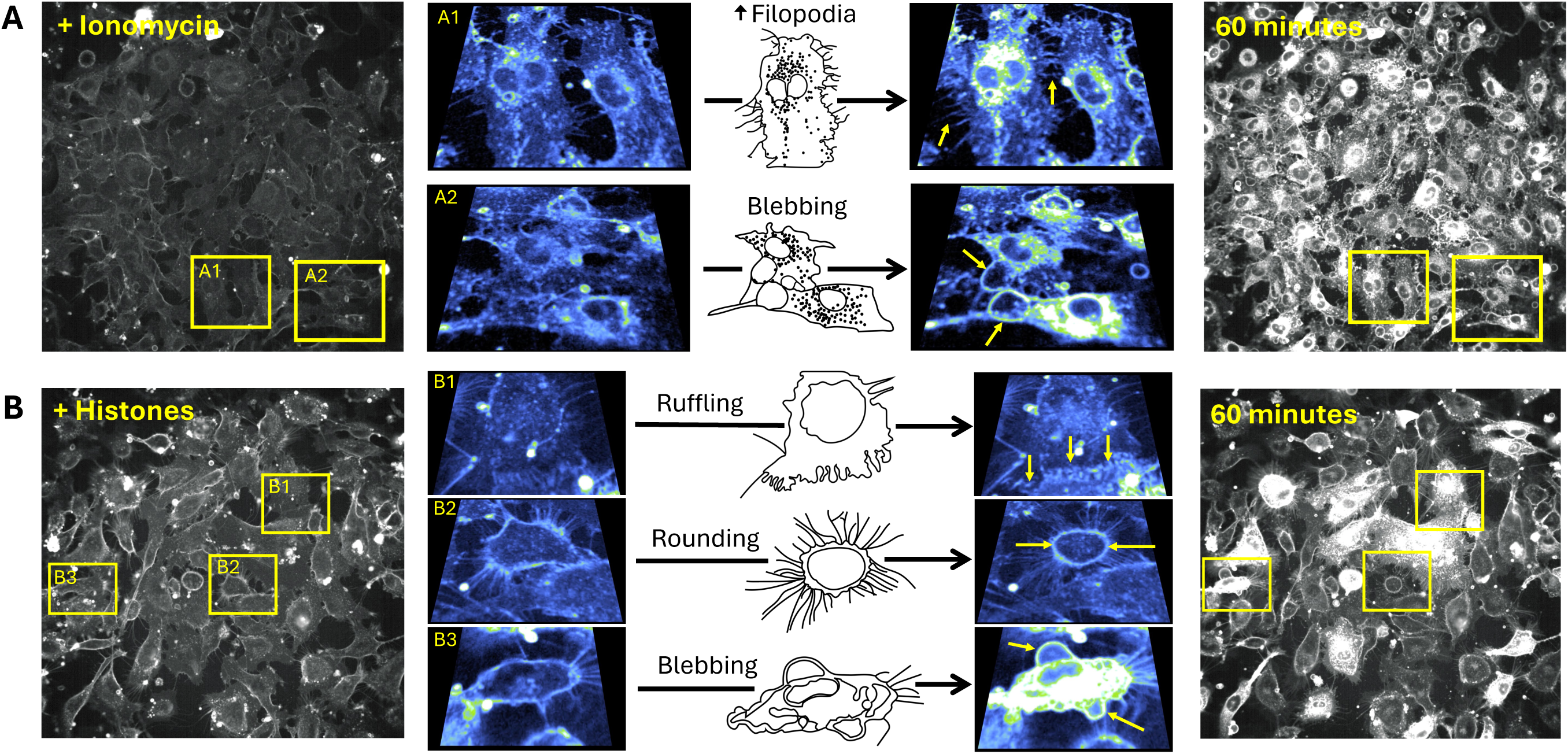
Histones and Ca²⁺ overload produce distinct patterns of plasma membrane movement. (**A**) During 60-min imaging of EC monolayers, ionomycin produced near-uniform responses across the field: rapid dye incorporation with punctate vesicular bodies, then elongating motile filopodia (A1; Supplemental Video 5) and terminal large blebs (A2; Supplemental Video 6). (**B**) Histones produced a heterogeneous, mosaic response in which some cells responded rapidly and others appeared unaffected (Supplemental Video 4). Dye uptake was patchy, often initiating at discrete regions before spreading. Unique to histones, membrane ruffling occurred at cell edges with undulating expansion of the membrane border (B1; Supplemental Video 7), while other cells underwent rapid rounding leaving retraction fibers (B2; Supplemental Video 8) or end-stage blebbing with extensive dye uptake (B3; Supplemental Video 9).

### Membrane injury is not caused by Ca²⁺ overload and is amplified by low extracellular Ca²⁺

Because histone-induced Ca²⁺ influx preceded membrane changes, we considered whether Ca²⁺ overload triggers the membrane disruption. If so, removing extracellular Ca²⁺, which prevents histone-induced Ca²⁺ entry,^7^ should prevent the membrane response. It did the opposite. In confluent monolayers (60±17 cells per field), histone exposure in 0 mM Ca²⁺ produced an exaggerated response: dye-permeable cells increased from 34±7% at 1.2 mM Ca²⁺ to 60±11%, and the mean maximum fluorescence change at 40 min rose from ΔF/F₀ of 6.56 to 18.7 (Figure 4A–C). Blebbing and deformation were correspondingly more extensive.

**Figure 4.**
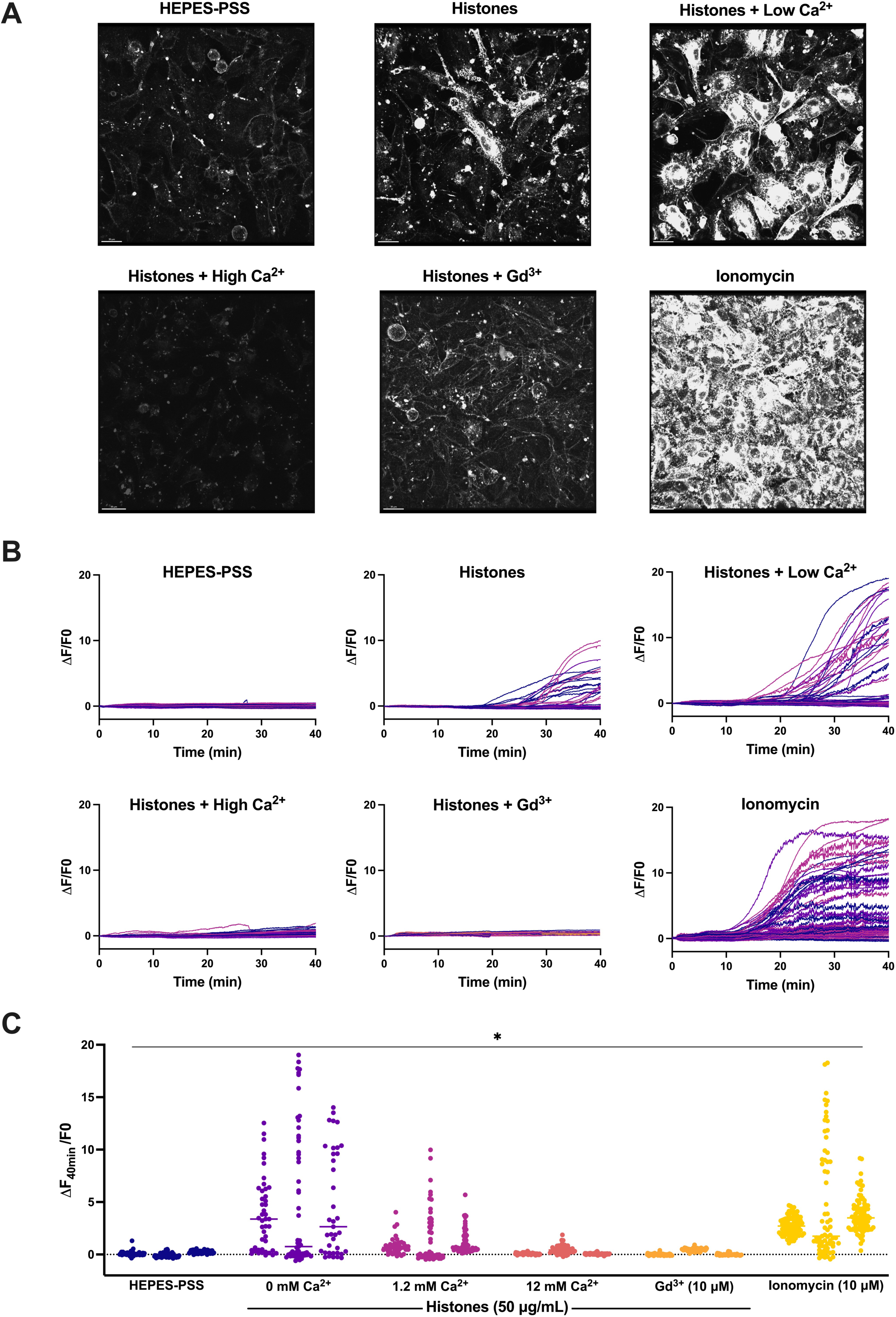
Extracellular Ca²⁺ sets the severity of histone-induced membrane injury. (**A**) Representative Z-stacks of cultured ECs after 40-min exposure to buffer alone (HEPES-PSS); histones (50 µg/mL) in low (0 mM), normal (1.2 mM), or high (12 mM) Ca²⁺; histones in normal Ca²⁺ after Gd³⁺ (10 µM) pre-treatment; or ionomycin (10 µM). (**B**) Representative traces of individual cell cytoplasmic fluorescence (ΔF/F₀) over 40 min for each condition. (**C**) Final fluorescence (ΔF₄₀ₘᵢₙ/F₀) for each cell with replicates displayed separately (n=3). Kruskal-Wallis test; P<0.05.

Barrier function showed the same dependence. Using electric cell-substrate impedance sensing, histones (100 µg/mL) in physiological Ca²⁺ initially *increased* normalized endpoint resistance and total resistance over one hour, consistent with an activation response. In low Ca²⁺, histones instead produced a near-immediate and significant fall in both normalized endpoint resistance (0.5±0.0, n=7 vs 1.7±0.1, n=8) and total resistance (AUC 462±9, n=7 vs 1558±81, n=8), beginning within 10 min (Supplemental Figure 3). Low extracellular Ca²⁺ therefore converts a transient barrier-stabilizing response into rapid barrier failure.

### Neutralizing membrane surface charge prevents histone-membrane interaction

We next tested whether high Ca²⁺ and Gd³⁺ protect live ECs. Both high Ca²⁺ (12 mM) and Gd³⁺ (10 µM) markedly diminished histone-induced membrane dye internalization by quantitative spatiotemporal analysis of ΔF/F₀ (Figure 4A–C). Flow cytometry confirmed this observation. Median FM1-43 fluorescence intensity increased 1.23-fold after histones (50 µg/mL) in 1.2 mM Ca²⁺ and 1.33-fold in 0 mM Ca²⁺, whereas cells stimulated in 12 mM Ca²⁺ showed a 1.36-fold *decrease* relative to unstimulated cells at the same Ca²⁺ concentration (Figure 5A).

**Figure 5.**
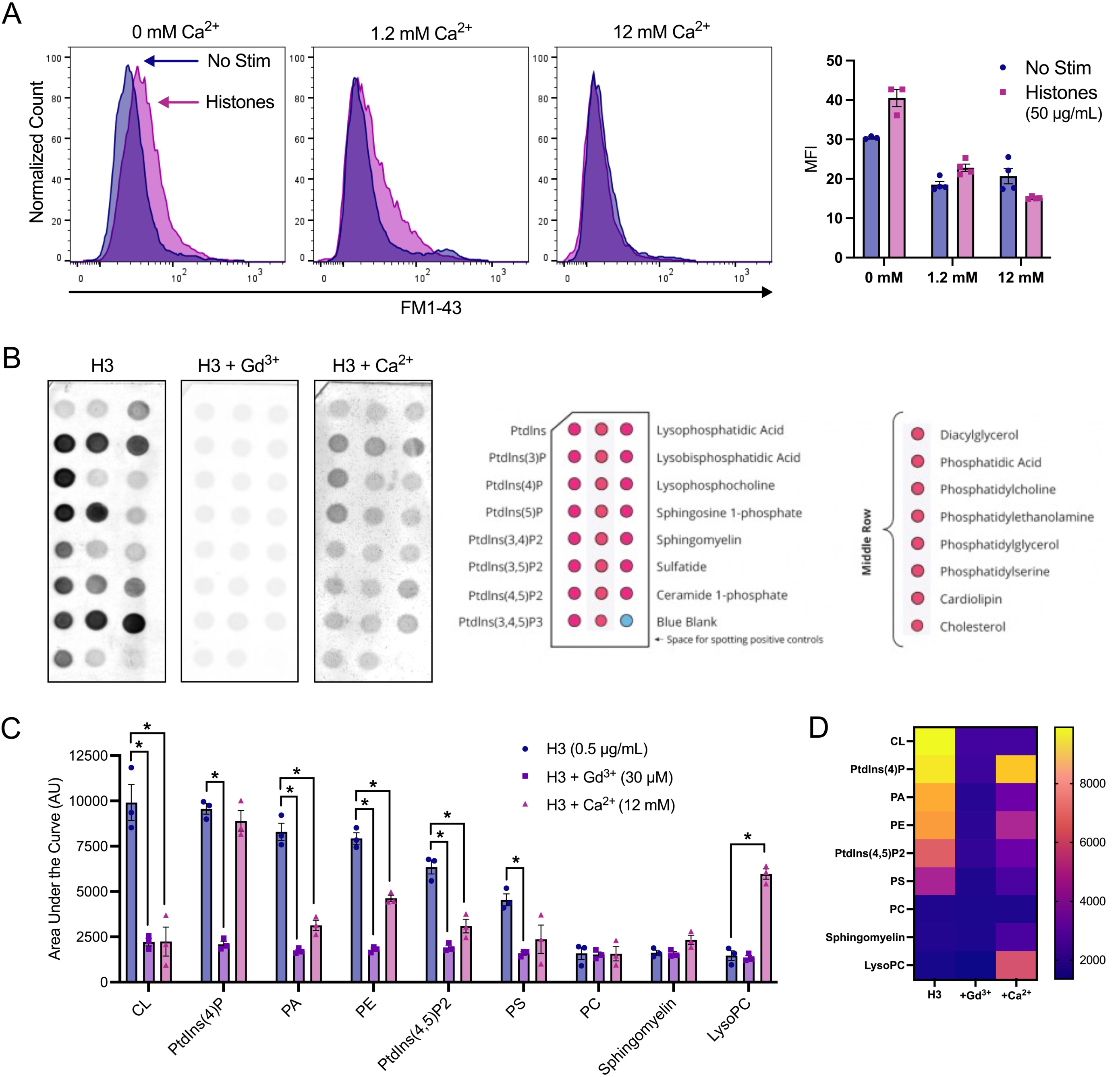
Charge screening by Ca²⁺ or Gd³⁺ prevents histone binding to anionic phospholipids. **(A**) Flow cytometry of FM1-43 fluorescence after incubation of ECs with histones (50 µg/mL) or ionomycin (10 µM) for 40 min in HEPES-PSS containing 0, 1.2, or 12 mM Ca²⁺. Representative histograms and median fluorescence intensity. (**B**) Representative lipid-binding strips exposed to histone H3-GST alone, with Gd³⁺ (30 µM), or with Ca²⁺ (12 mM) (left), with schematic of lipid identities adapted from the manufacturer (right). (**C**) Summary data for histone H3 binding to nine of the phospholipids on the strips shown in (B), alone or with Gd³⁺ (30 µM) or Ca²⁺ (12 mM); n=3 per group. (**D**) Heat map of the same data. Quantification and statistics in Supplemental Methods.

To test the electrostatic mechanism directly, we measured binding of glutathione S-transferase (GST)-tagged histone H3 to individual membrane phospholipids. The histone proteins bound anionic but not zwitterionic phospholipids in the absence of Ca²⁺, consistent with an electrostatic interaction shielded by Ca²⁺ (Figure 5B). Gd³⁺ (30 µM) completely abolished histone-phospholipid binding, removing all preferential binding, and high Ca²⁺ (12 mM) significantly reduced binding to most phospholipids (Figure 5C, D). Notably, micromolar Gd³⁺ prevented histone binding, whereas millimolar Ca²⁺ was required for the same block, consistent with the greater charge and charge density of Gd³⁺.

### Phosphatidylserine externalization occurs without extracellular Ca²⁺

Histone-induced endothelial Ca²⁺ signals may contribute to thrombosis in part by translocating phosphatidylserine (PS) to the membrane surface, either through the Ca²⁺-activated scramblases TMEM16E and F or through cell death.^38–41^ We asked whether histone-induced PS externalization is modulated by extracellular Ca²⁺ and how it relates to permeability to propidium iodide (PI). After 1 h of histone exposure in 0, 1.2, or 12 mM Ca²⁺, cells were stained with Annexin V and PI (Figure 6).

**Figure 6.**
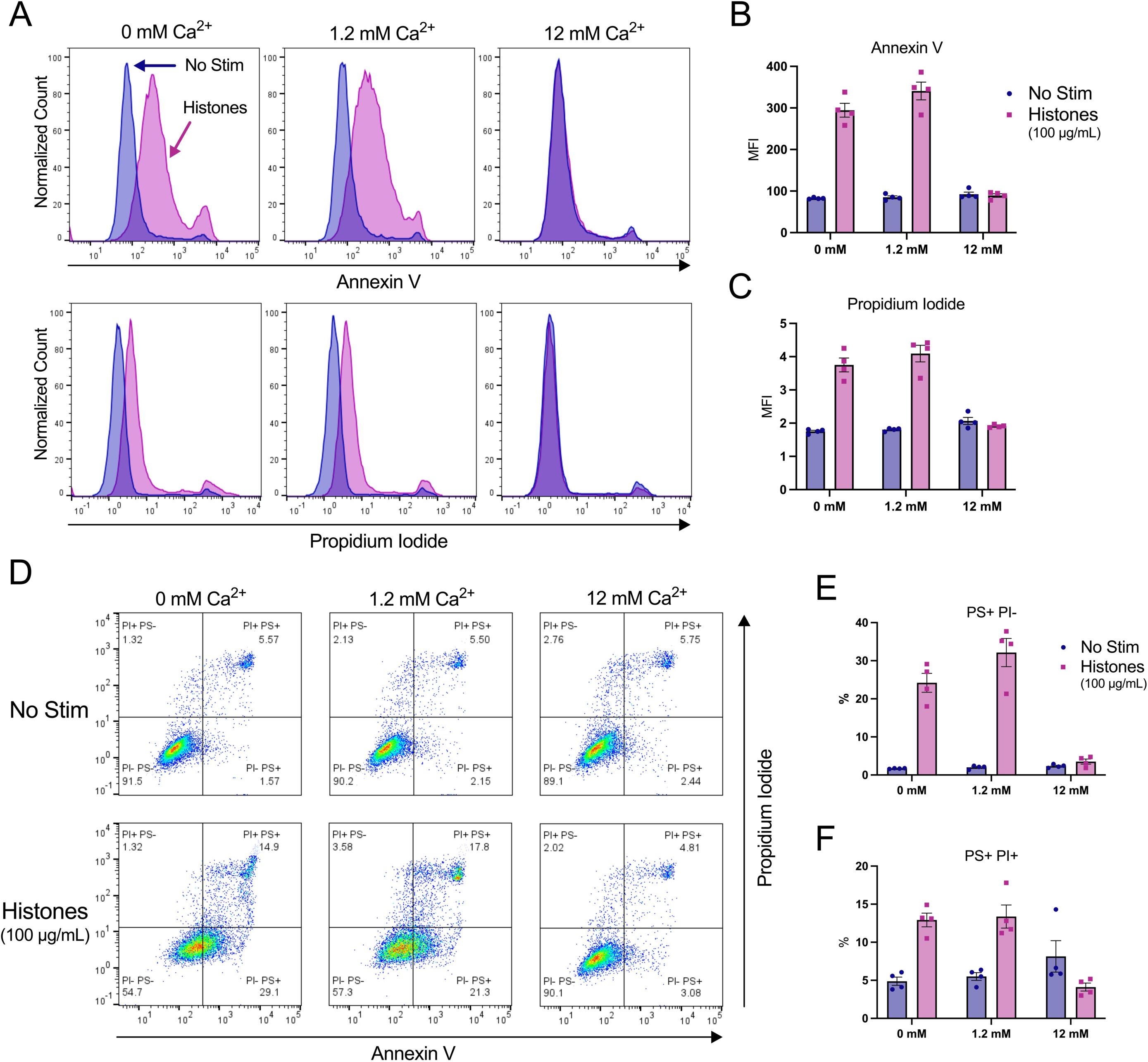
High-dose histones externalize phosphatidylserine and permeabilize ECs independently of extracellular Ca²⁺. ECs treated with histones (100 µg/mL) in HEPES-PSS containing 0, 1.2, or 12 mM Ca²⁺ for 1 h. Following histone exposure, all samples were transferred to the same Annexin V binding buffer, independent of the Ca²⁺ concentration during treatment, and stained with Annexin V and propidium iodide. (**A**) Representative histograms of Annexin V (top) and PI (bottom) fluorescence for buffer alone (blue) or histones (pink). (**B, C**) Median fluorescence intensity for Annexin V and PI (n=4). (**D**) Representative dot plots with gating strategy. (**E, F**) Percent PS⁺/PI⁻ and PS⁺/PI⁺ cells.

At 100 µg/mL, a dose we have previously shown produces PS translocation,^41^ 32.15±7.37% of cells were PS⁺/PI⁻ at 1.2 mM Ca²⁺, 24.20±4.95% at 0 mM, and 3.52±1.32% at 12 mM, representing 16-, 14-, and 1.5-fold increases over unstimulated controls at the corresponding Ca²⁺ concentration. Double-positive (PS⁺/PI⁺) fractions were 13.38±3.02% at 1.2 mM, 12.93±1.82% at 0 mM, and 4.11±1.05% at 12 mM (Figure 6D–F). Substantial PS externalization therefore occurs even in the absence of extracellular Ca²⁺, demonstrating that PS exposure does not require extracellular Ca²⁺ influx, although a contribution from intracellular Ca²⁺ stores or Ca²⁺-independent mechanisms cannot be excluded. One caveat applies to the protection by high Ca²⁺: unstimulated cells in 12 mM Ca²⁺ already showed elevated PS⁺/PI⁺ staining (8.14±4.13% vs 5.51±1.01% in normal Ca²⁺), so the apparent protection at 12 mM is measured against a raised baseline. At 50 µg/mL histones, the same Ca²⁺ dependence was present but smaller in magnitude (Supplemental Figure 2).

### Histone-induced membrane permeabilization is reparable in a subset of endothelial cells

High-dose histones kill ECs^3,7^, but the heterogeneity of the response across cells within a field raised the possibility that some permeabilized cells recover. No existing assay distinguishes a cell that was permeabilized and resealed from one that was never permeabilized, so we developed a sequential two-dye live-cell strategy to measure membrane recovery (Supplemental Figure 4). Live ECs were stained with Hoechst and SYTOX Green to establish baseline viability, washed, and then exposed for 1 h to buffer, histones (100 µg/mL), or ionomycin (10 µM). SYTOX Green was reapplied and cells imaged to identify those permeabilized during stimulation. Cells were then returned to culture conditions for 3 h and stained with spectrally distinct SYTOX AADvanced. This design resolves three phenotypes: *intact* (negative for both dyes), *sustained permeability* (positive for both), and *repaired* (SYTOX Green-positive but AADvanced-negative cells). In other words, a repaired cell appears green, because it took up the green dye from its initial wound, repaired its plasma membrane, and subsequently excludes the red dye. Surprisingly, we observed a discrete population of green cells that excluded red dye entry, indicating successful plasma membrane repair is feasible after injury by histones (Figure 7A). Critically, ionomycin-treated cells showed no repaired population in any trial, establishing that the phenotype is not an artifact of sequential staining; these Ca²⁺-overloaded cells do not reseal. The size of the repaired population varied with the recovery medium (Figure 7B).

**Figure 7.**
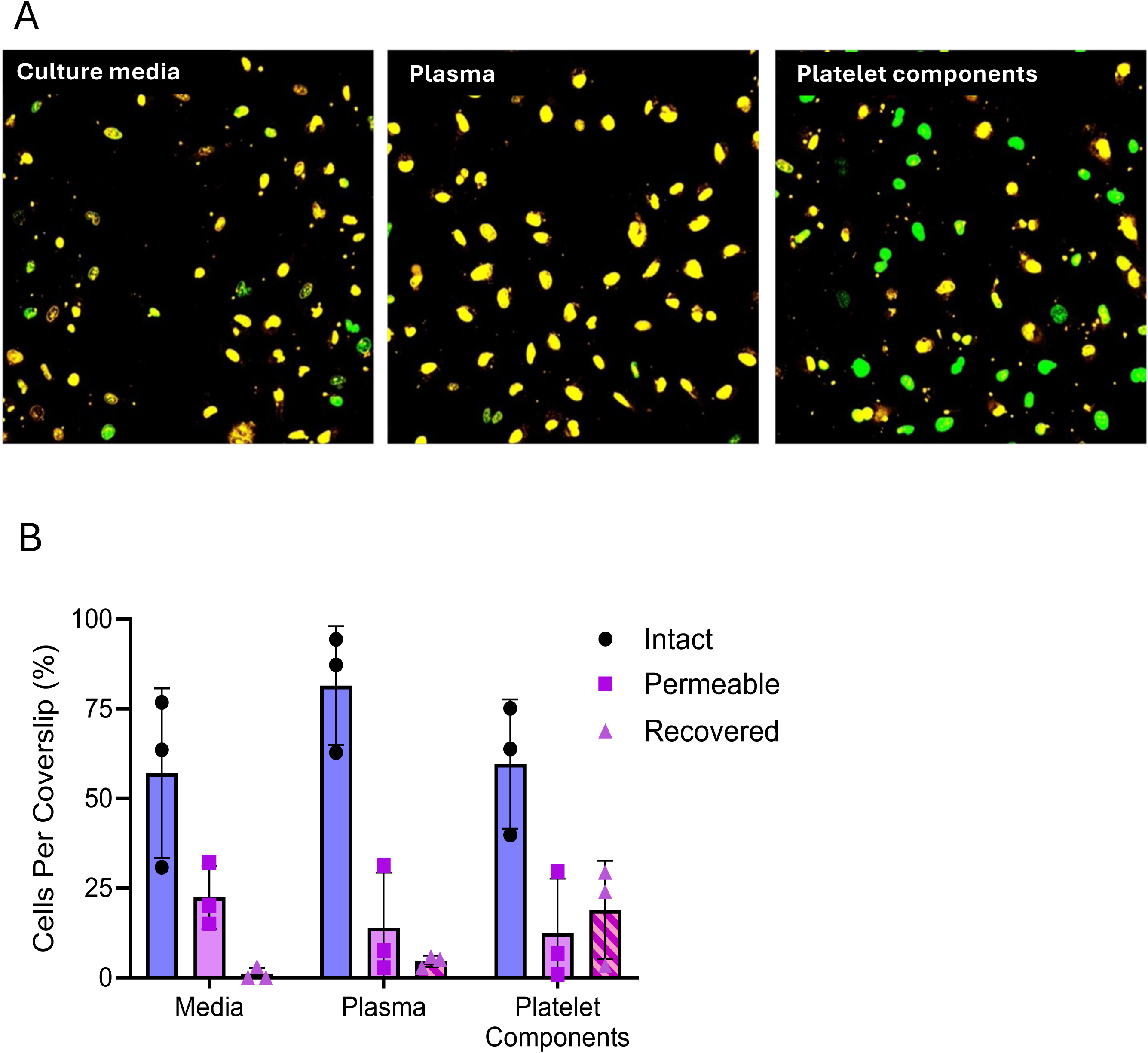
Histone-induced membrane permeabilization is reparable in a subset of endothelial cells. ECs were injured for 1 h with histones (100 µg/mL), then allowed 3 h of recovery in culture medium, healthy-donor human plasma, or healthy-donor platelet components. Sequential staining with SYTOX Green before recovery and SYTOX AADvanced after recovery resolves intact, sustained-permeability (labeled Permeable) and repaired (labeled Recovered) phenotypes; the assay design is shown in Supplemental Figure 4. (**A**) Representative merged images after recovery in each medium. (**B**) Percent of cells per coverslip in each phenotype; each point is the mean of three fields within one coverslip, bars show mean ± SD. Differences in the repaired fraction between recovery media were not significant (medium versus platelet components P=0.299, medium versus plasma P=0.951). Ionomycin-injured cells showed no repaired population in any trial.

We then asked whether the composition of the extracellular milieu during recovery influences repair, comparing culture medium and human blood components as recovery media.

These were platelet components, collected from isolated platelets following release of their contents,^42^ and human plasma from the same individual donors. The fractions of recovered endothelial cells were 1.0±1.73% in medium, 4.6±1.6% in plasma, and 18.9±13.7% in platelet components (Figure 7B). This ordering is consistent with a milieu-dependent contribution to repair, but the differences did not reach statistical significance by two-way ANOVA with Tukey correction (medium versus platelet components, P=0.299; medium versus plasma, P=0.951), and the variance in that condition was large relative to its mean. We therefore regard the identity of the repair-promoting constituents, the magnitude of any milieu effect, and heterogeneity across platelet donors as unresolved, requiring a study poweredfor this comparison.

One candidate route by which extracellular constituents could act is direct vesicle delivery to the injured membrane. Live membrane imaging showed that individual extracellular vesicles approached the endothelial surface and tethered at the cell surface, initiating a wave of membrane activity that propagated along the cell at approximately 10 µm/min (Figure 8A–C). Following attachment, vesicles remained at the site and endocytosis began within approximately one minute, with membrane fluorescence accumulating at the site for 40 min (Figure 8D). In some recordings, vesicles released by one cell were taken up by adjacent and remote cells and passed along connected lamellipodia and filopodia (Supplemental Video 12). This resembled the “exosome surfing” described previously, in which filopodia capture single vesicles and convey them to endocytic hot spots at the filopodial base.^43^ Comparable behavior was observed in surgically opened *en face* vessels after mechanical injury with microscissors. Vesicles moved freely above the endothelial surface and tethered preferentially to cells at the injured margin, while uninjured cells in the same field acquired few vesicles, and the free vesicle population declined over the recording (Supplemental Figure 5A–E).

**Figure 8.**
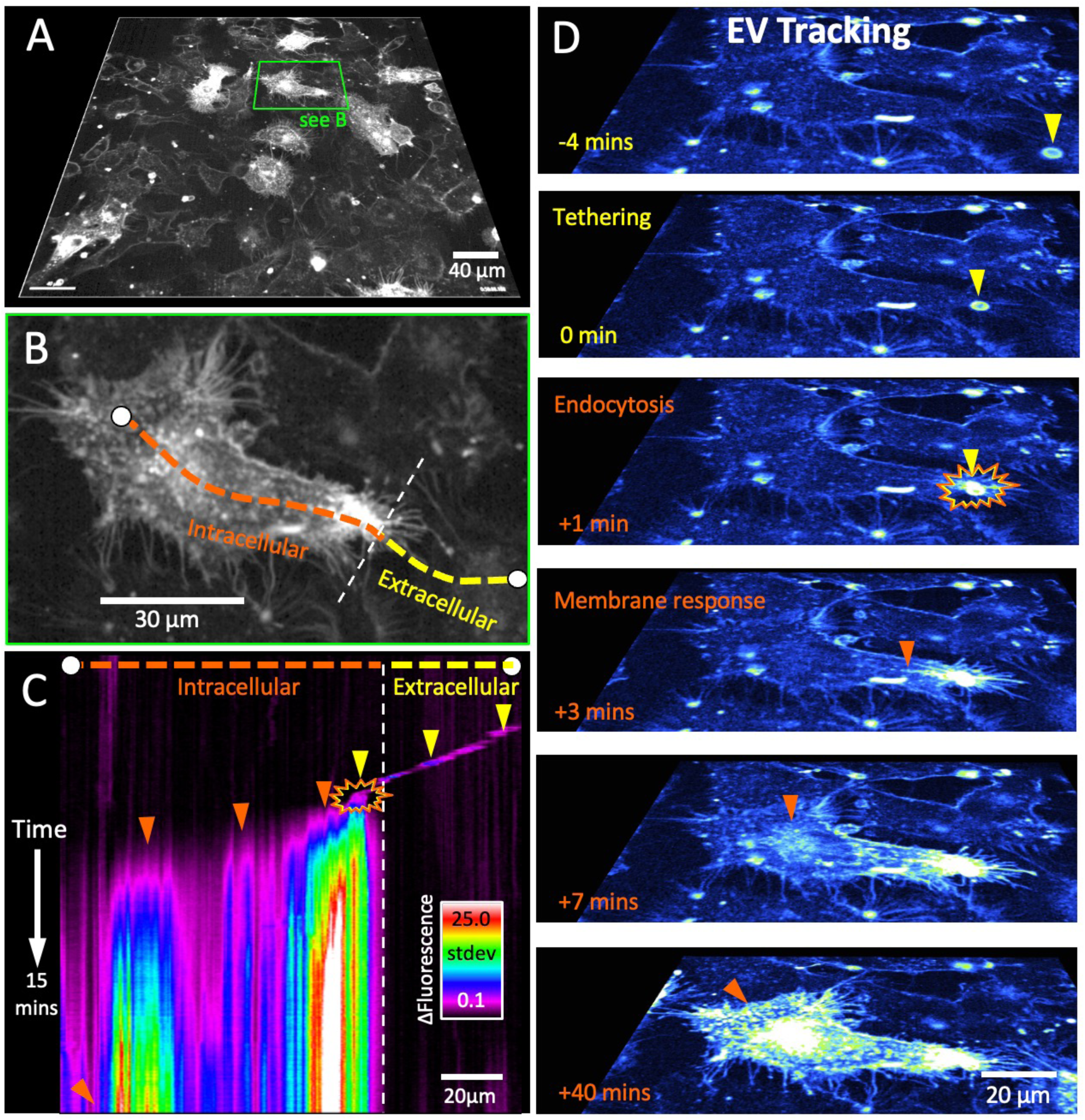
Endothelial cells tether and internalize extracellular vesicles within minutes of contact. (**A**) Planar view of an FM1-43-stained EC monolayer, showing (**B**) the cell and spline used to generate the spatiotemporal map of membrane fluorescence. (**C**) Spatiotemporal map showing the trajectory of an extracellular vesicle (yellow arrowheads) before attachment to the EC, generating a wave of membrane activity propagating from the attachment point at approximately 10 µm/min (orange arrowheads). (**D**) Vesicle approaches the cell tip and attaches (yellow arrowheads); endocytosis begins within one minute (orange), with accumulated fluorescence signal over the subsequent 40 min shown in the bottom panel.

Extracellular vesicle-enriched preparations from stimulated ECs contained predominantly CD63-positive particles, with smaller CD9-positive and CD81-positive populations (Supplemental Figure 5F, G). The tetraspanin profile depended on the agonist: particles released following histone stimulation showed greater CD63 and lower CD81 labeling than those released following ionomycin stimulation, resembling the profile observed in platelet-derived preparations. These observations establish that ECs both release and internalize extracellular vesicles during membrane injury; they do not establish that vesicle uptake mediates the repair we measured, and we have not yet tested vesicle-depleted preparations.

## Discussion

Cardiovascular shock from trauma and sepsis is among the leading causes of morbidity and mortality worldwide.^44,45^ Recognition of endotheliopathy as a driver of coagulopathy and poor outcome in shock was an important advance, and the shock-induced endotheliopathy paradigm explains how different shock states converge on similar patterns of endothelial activation and injury.^2^ Here we define a mechanism for histone-induced endothelial injury in which extracellular Ca²⁺ acts not as the agent of toxicity but as its principal restraint, and we show that the resulting membrane injury is not uniformly irreparable.

We report five findings. First, store-operated Ca²⁺ entry (SOCE) does not contribute to histone-induced cytosolic Ca²⁺ influx: deletion of plasma membrane ORAI1/2/3 channels had no effect, and the Gd³⁺ block required concentrations two orders of magnitude above the SOCE IC₅₀. Second, histones bind and induce deformations of endothelial membranes. Third, Ca²⁺ influx is not required for histone-induced membrane deformation. Fourth, histone-induced membrane permeabilization depends on electrostatic interaction with anionic phospholipids: it is exacerbated by low Ca²⁺ and blocked by mM Ca²⁺ or µM Gd³⁺. Fifth, we demonstrate that histone-induced permeabilization is reparable. Separately, the Ca²⁺ transients evoked by histones were irregular, and distinct both from the regenerative oscillations evoked by physiological agonists and from the sustained plateau produced by ionomycin.

The inverse relationship between extracellular Ca²⁺ and injury severity is the central mechanistic result and was unexpected. Because removing extracellular Ca²⁺ prevents histone-induced Ca²⁺ entry,^7^ Ca²⁺ overload was the natural candidate trigger for membrane disruption. Instead, Ca²⁺ removal amplified dye uptake, deformation, and barrier failure, while elevated Ca²⁺ or Gd³⁺ abolished histone binding to anionic phospholipids. The coherent interpretation is a surface charge mechanism. The plasma membrane is more electronegative than organellar membranes,^46^ and in ECs the glycocalyx confers additional net negative charge.^47^ This electronegative field is physiologically protective, repelling circulating platelets and anionic blood components, but it also recruits extrinsic cationic proteins such as histones. Divalent and trivalent cations screen that field: the same surface charge theory invoked to explain membrane stabilization by Ca²⁺ in nerve and cardiac muscle^25,26^ accounts for the protection we observe, and the far lower concentration of Gd³⁺ required (µM versus mM) follows from its greater charge and charge density. Membrane-bending by surface-bound proteins, which produces invaginations and vesiculation,^48^ may further explain how histone binding progresses from binding to deformation and vesicle release.

The physiological importance of endothelial surface electronegativity suggests this vulnerability is a trade-off rather than a design flaw. Maintaining the non-thrombogenic state of the intima requires a field that necessarily also recruits extrinsic cationic proteins. Extracellular histones can therefore be understood as exploiting an electrostatic feature that exists for an unrelated protective purpose, which is why the degree of screening by ambient Ca²⁺ (which is lost during citrate-based resuscitation) becomes a determinant of injury.

This mechanism reframes hypocalcemia during resuscitation. Low ionized Ca²⁺ predicts transfusion requirement and mortality in trauma,^10,12^ and massive transfusion protocols diverge on whether to give Ca²⁺ routinely or only as needed.^9,11,13^ That debate has centered on coagulation-factor function. Our data identify a distinct, parallel liability of hypocalcemia: at low ionized Ca²⁺, the endothelial surface is less screened and more vulnerable to the cationic histones that circulate at high concentration in these patients. If this holds *in vivo*, hypocalcemia is not only a coagulation problem but a determinant of endothelial susceptibility, and the relevant clinical question becomes whether earlier Ca²⁺ correction protects the endothelium and not just clotting enzymes.

The heterogeneity of the endothelial response was striking and reproducible. Only a subset of cells responded to histones, and even within responding cells membrane changes began at discrete hot spots before spreading. The origin of those hot spots may be extrinsic rather than cell-intrinsic: vesicle contact preceded dye uptake at the same site (Figure 8). We recently observed a comparable selectivity following focal laser injury of endothelial monolayers: vesicles released from a destroyed cell are progressively captured by surviving injured neighbors as those cells externalize phosphatidylserine, while vesicles are not efficiently captured by cells that are uninjured.^49^ Vesicle capture therefore appears to track the injury state of the receiving membrane rather than its activation state. Because histones externalize phosphatidylserine in a subset of endothelial cells (Figure 6), the same injury that permeabilizes the membrane may also generate the anionic surface that recruits vesicles to it. Distinct subpopulations may act as sentinel cells with heightened susceptibility to chemical or mechanical insult, amplifying danger signals to neighbors. Non-sentinel cells may respond by protruding thin ruffling lamellipodia that transiently increase barrier function, which would explain the early rise in impedance we and others observe before barrier degradation over 12–24 h.^39,50^ PS externalization may reinforce this heterogeneity as a feed-forward loop: PS is strongly anionic and avidly binds histones (Figure 5), so activated or pre-apoptotic cells displaying surface PS should become preferentially susceptible to further histone binding. Moreover, PS-displaying extracellular vesicles may aggregate free histone proteins and deliver them to cells.

Direct electrostatic interaction with the bilayer is unlikely to be the only route by which histones act on cells, and our data do not exclude others. Histones also engage Toll-like receptors 2 and 4, the purinergic receptor P2X7, and the receptor for advanced glycation end-products, and downstream inflammatory and pyroptotic responses vary with histone post-translational modification, subunit composition, DNA content, and cell type.^23,38,51–59^ This multiplicity may explain why sensitivity to histones differs so markedly between cell types. Neurons are relatively resistant compared with macrophages or ECs, for example.^60^ A plausible synthesis is that direct bilayer disruption sets the physical injury, measured here, while receptor-mediated signaling shapes the inflammatory response that follows. Whether charge screening modifies only the bilayer disruption or both this and the inflammatory response remains to be tested, as does whether the histone sensitivity of other vascular cells (smooth muscle, erythrocytes, platelets) is similarly Ca²⁺-dependent.^38,52,58^

Our most translationally consequential observation is that histone-induced permeabilization of the endothelial plasma membrane bilayer is reparable. The extent to which acutely injured endothelium recovers through cellular repair, angiogenesis, or recruitment of circulating progenitors is poorly understood.^61–64^ By staining sequentially with two spectrally distinct viability dyes, we identified cells that were demonstrably permeabilized during injury and subsequently prevented entry of a secondary dye; this shows EC plasma membranes have the capacity to reseal after a thromboinflammatory insult. Our observation that ionomycin-treated cells never showed this phenotype supports the interpretation that the assay detects restoration of membrane impermeability, and that the mode of injury determines reparability: graded, electrostatically driven lesions appear reparable, whereas global Ca²⁺ overload does not. We also observed heterogeneity in recovery, suggesting functionally distinct subpopulations of ECs, such that histones may cause irreparable damage to highly susceptible cells and transient, reparable injury to others.

Whether the extracellular milieu determines repair efficiency remains an open question. Repaired fractions trended higher in plasma and higher still in platelet components than in culture medium, consistent with the hypothesis that donor blood products contain constituents that support endothelial membrane repair, and with clinical observations that early plasma transfusion lowers biomarkers of endothelial injury.^14^ But this comparison was not powered and did not reach significance, and donor-to-donor variability in platelet component composition is a plausible source of its large variance. Such variability is unlikely to be incidental: longitudinal multiomic profiling of injured patients resolves distinct thromboinflammatory endotypes and trajectories,^65^ implying that both the injurious and the reparative constituents of plasma differ substantially between individuals. We also found that ECs tether and internalize extracellular vesicles within minutes of contact, which offers a mechanistically attractive candidate for membrane delivery: platelets and platelet vesicles can plug and repair endothelial lesions^66^ and prevent thrombin-induced barrier breakdown.^42,67,68^ Yet, we did not test vesicle-depleted preparations and cannot attribute the repair we measured to vesicles. Resolving this requires defined fractionation of donor material into vesicle and non-vesicle components, dosing by particle number rather than dilution, with sufficient power to treat donor heterogeneity in composition as a covariate.

Several limitations bound these conclusions. Excluding store-operated entry does not identify the route by which Ca²⁺ does enter, and reverse-mode Na⁺/Ca²⁺ exchange, which can operate when cells depolarize, was not tested here. That pathway remains open, although its identity is secondary to the present conclusions: membrane deformation and permeabilization proceeded in nominally Ca²⁺-free solution and were amplified rather than prevented by removing extracellular Ca²⁺. Several experiments used immortalized cell lines and ex vivo arterial preparations in buffers with Ca²⁺ concentrations exceeding the physiological range, so the magnitude of effects *in vivo* in the presence of albumin and other plasma proteins remains to be established. Gd³⁺ is pharmacologically non-specific; we use it as a surface charge probe rather than a candidate therapeutic, and the concentration required for lipid-strip blockade (30 µM) exceeded that used in cells (10 µM). The apparent protection by 12 mM Ca²⁺ in PS assays is measured against an elevated baseline. Several mechanistic panels rest on small numbers of independent replicates. The sentinel-cell interpretation is inferred from response distributions rather than demonstrated by prospective identification of susceptible cells. Finally, the clinical inference connecting hypocalcemia to endotheliopathy in patients is extrapolated from *in vitro* electrostatics and requires testing against measured ionized Ca²⁺ and circulating histone levels in injured patients.

Within those bounds, two robust conclusions emerge. Extracellular Ca²⁺ limits histone-induced endothelial membrane injury by electrostatic screening rather than by supplying Ca²⁺ influx, so hypocalcemia is a plausible direct amplifier of shock-induced endotheliopathy. Also, endothelial membrane repair after histone injury is a real, measurable process confined to a subpopulation of cells. This suggests that modifying the determinants of that repair, including donor plasma composition, is a tractable target for intervention in acute endotheliopathy.

## Acknowledgments

We thank Nicole Bouffard and Douglas Taatjes for guidance and support with ECIS measurements, performed at the Microscopy Imaging Center at the University of Vermont (RRID# SCR_018821).

## Sources of Funding

Research reported in this publication was supported by the National Institute of General Medical Sciences of the National Institutes of Health under award number R35GM144099 and its equipment supplement R35GM144099-02S3, and by the National Heart, Lung, and Blood Institute under award number R01HL166944. Together these awards supported the majority of this work, more than 50 percent: R01HL166944 supported the experiments shown in Figures 1 and 7, and R35GM144099 with its equipment supplement supported all other figures. Additional awards provided key resources, approaches and expertise: the National Institute of General Medical Sciences under award numbers P20GM125498 and P20GM135007 (Core C: Customized Physiology and Imaging Core); the National Heart, Lung, and Blood Institute under award number R35HL150778; the National Institute of Neurological Disorders and Stroke and the National Institute on Aging under award numbers RF1NS128963, R01NS110656 and R01NS119971; and the Office of the Director of the National Institutes of Health under award number S10OD026843, which supported shared imaging instrumentation. The total costs of the primary awards supporting this work were $2,200,000 for R35GM144099 and $12,130,497 for R01HL166944, each financed 100 percent with federal funds. Non-governmental support also contributed to this work in forms that are not allocated to individual projects and cannot be quantified, including support from the Totman Medical Research Trust, institutional support of investigator effort such as salary above the federal cap, and institutional matching funds for shared imaging instrumentation. The additional awards listed above contributed principally investigator effort and access to shared resources, which likewise cannot be apportioned precisely to this publication. The content is solely the responsibility of the authors and does not necessarily represent the official views of the National Institutes of Health.

## Disclosures

None.

## Data Availability

All imaging data, videos, and source data are available from the corresponding author on request.

K.F. had full access to all the data in the study and takes responsibility for its integrity and the data analysis.

## Non-standard Abbreviations and Acronyms

EC: endothelial cell
ECIS: electric cell-substrate impedance sensing
EV: extracellular vesicle
GST: glutathione S-transferase
MFI: median fluorescence intensity
PI: propidium iodide
PS: phosphatidylserine
SOCE: store-operated Ca²⁺ entry
TKO: triple knockout.

## Novelty and Significance

### What Is Known

- Extracellular histones released from injured cells and neutrophil extracellular traps in trauma and sepsis drive endothelial injury, coagulopathy, and organ failure.
- Histones raise endothelial cytosolic Ca²⁺ and degrade endothelial barrier function, but the route of Ca²⁺ entry has not been identified.
- Hypocalcemia is common during resuscitation with citrated blood products and is associated with increased transfusion requirement and mortality, an association generally attributed to impaired coagulation-factor function.

### What New Information Does This Article Contribute

- Histone-induced endothelial Ca²⁺ entry does not require store-operated Ca²⁺ entry (SOCE); genetic deletion of ORAI1/2/3 has no effect, and Gd³⁺ blocks this Ca²⁺ entry only at concentrations that screen membrane surface charge.
- Extracellular Ca²⁺ sets the severity of histone-induced membrane injury by charge screening rather than by supplying Ca²⁺ influx: lowering extracellular Ca²⁺ amplifies membrane deformation, permeabilization, and barrier failure, whereas elevated Ca²⁺ or µM Gd³⁺ prevents histone-phospholipid binding and protects cells.
- Histone-induced membrane permeabilization is not uniformly lethal. Using sequential live-cell viability imaging, we show that a subset of permeabilized endothelial cells recover membrane integrity within hours, establishing endothelial membrane repair as a measurable and potentially targetable process after thromboinflammatory injury.

## Notes

### Competing Interest Statement

The authors have declared no competing interest.

### Summary of Updates

Major revision including new results and figures

