## Supplemental material for "Charge Screening by Extracellular Ca²⁺ Shields Endothelial Cells from Histone Injury and Permits Membrane Repair"

Supplemental Figure 1

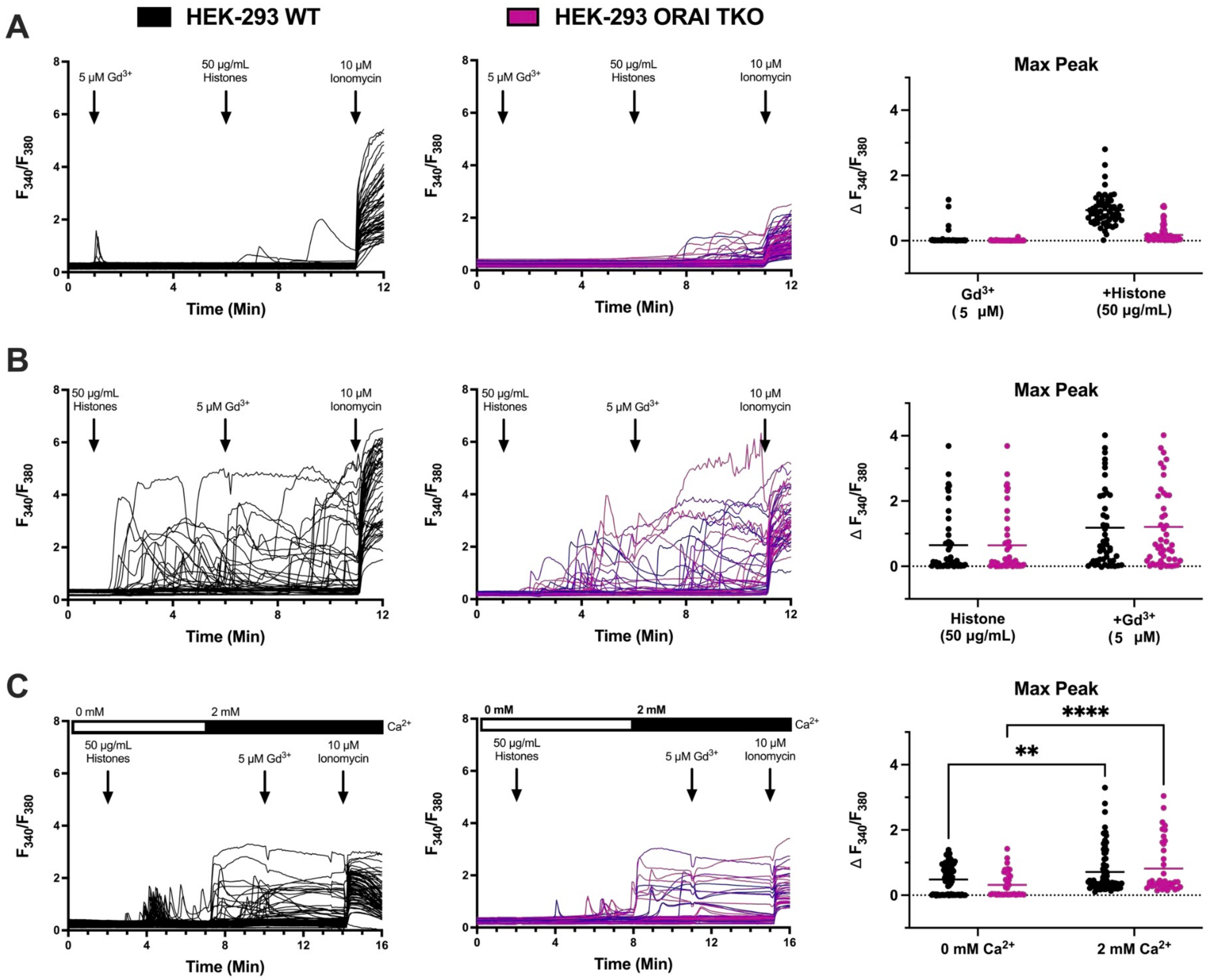

### Supplemental Figure 2

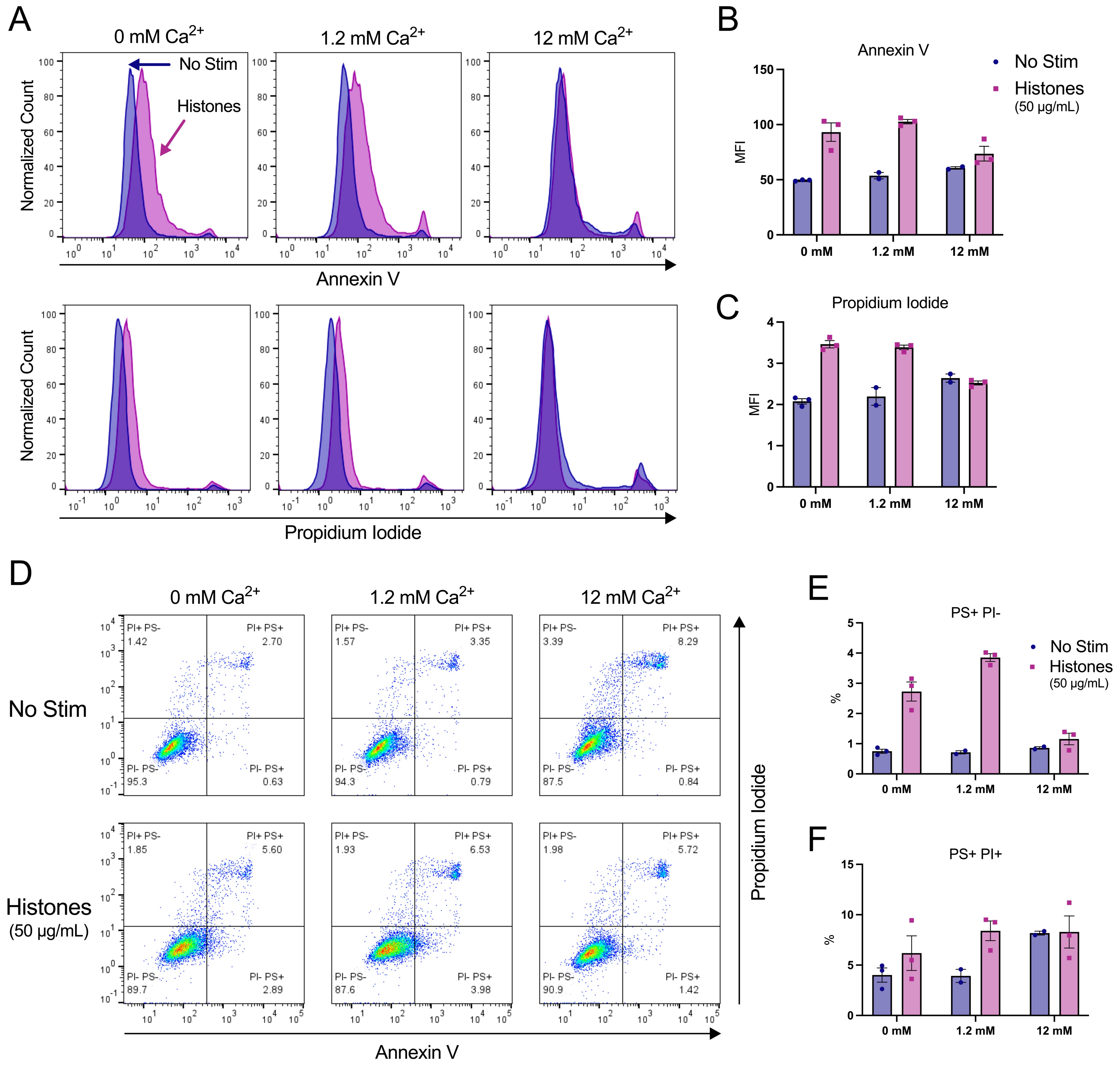

Supplemental Figure 3

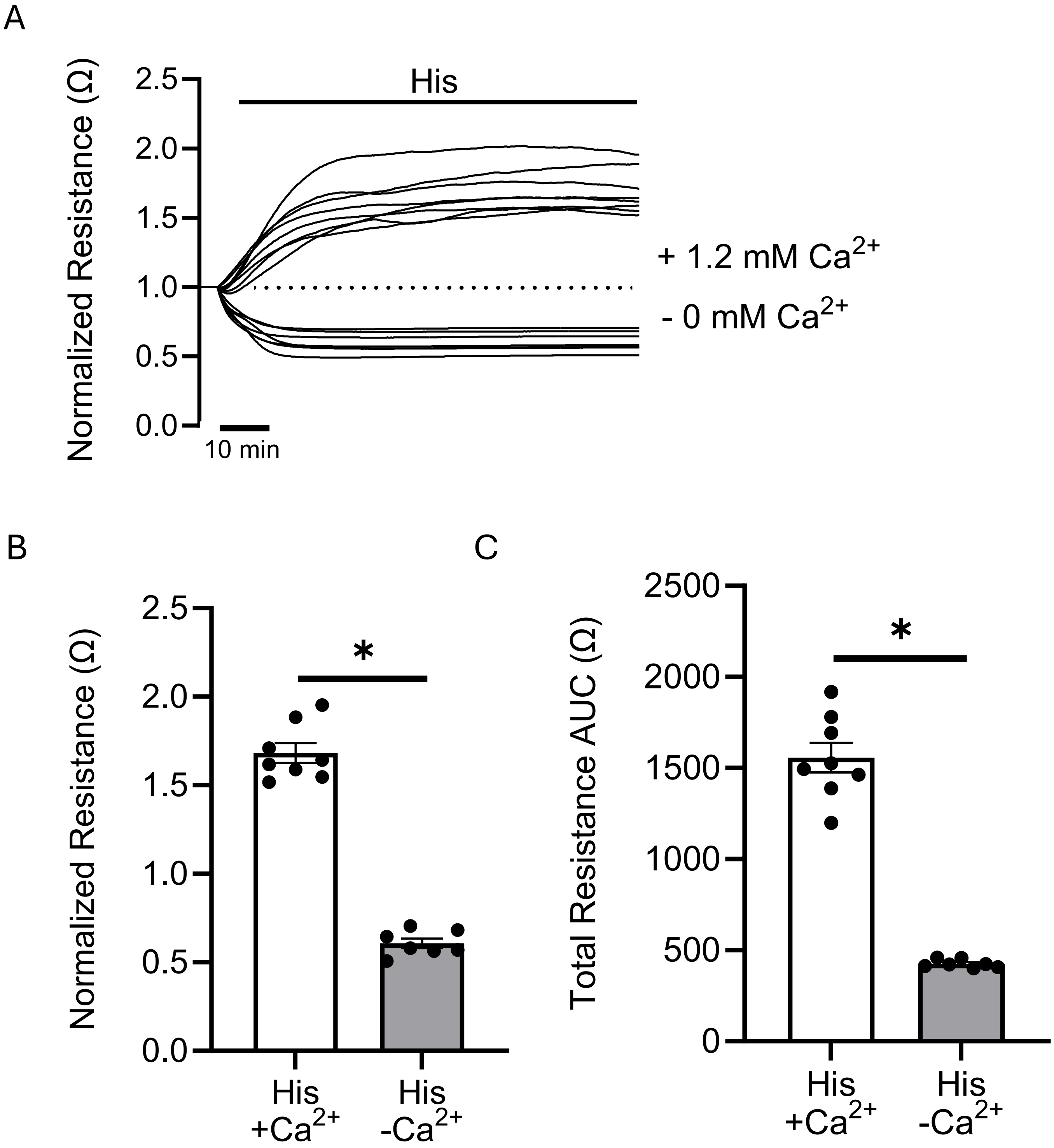

Supplemental Figure 4

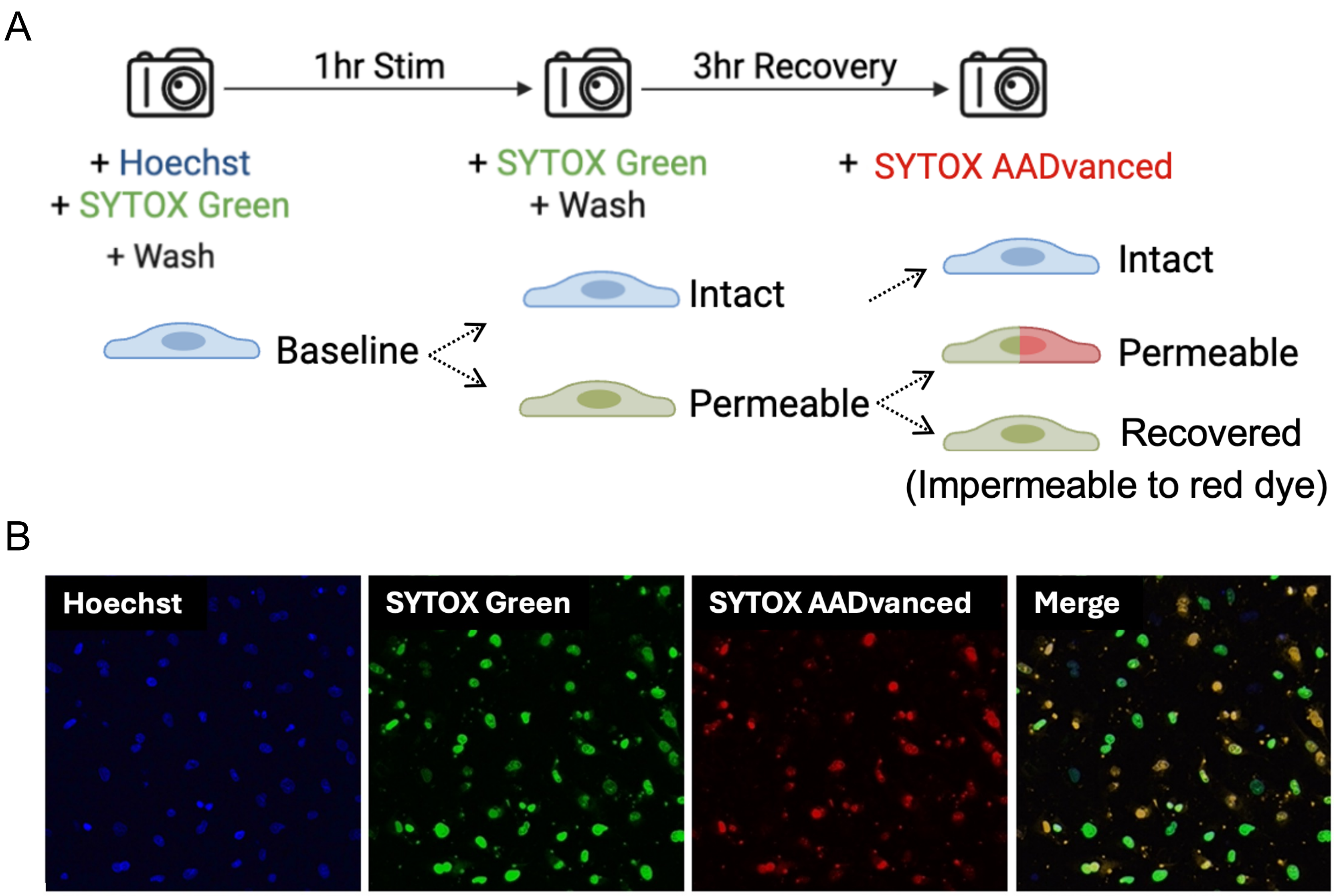

Supplemental Figure 5

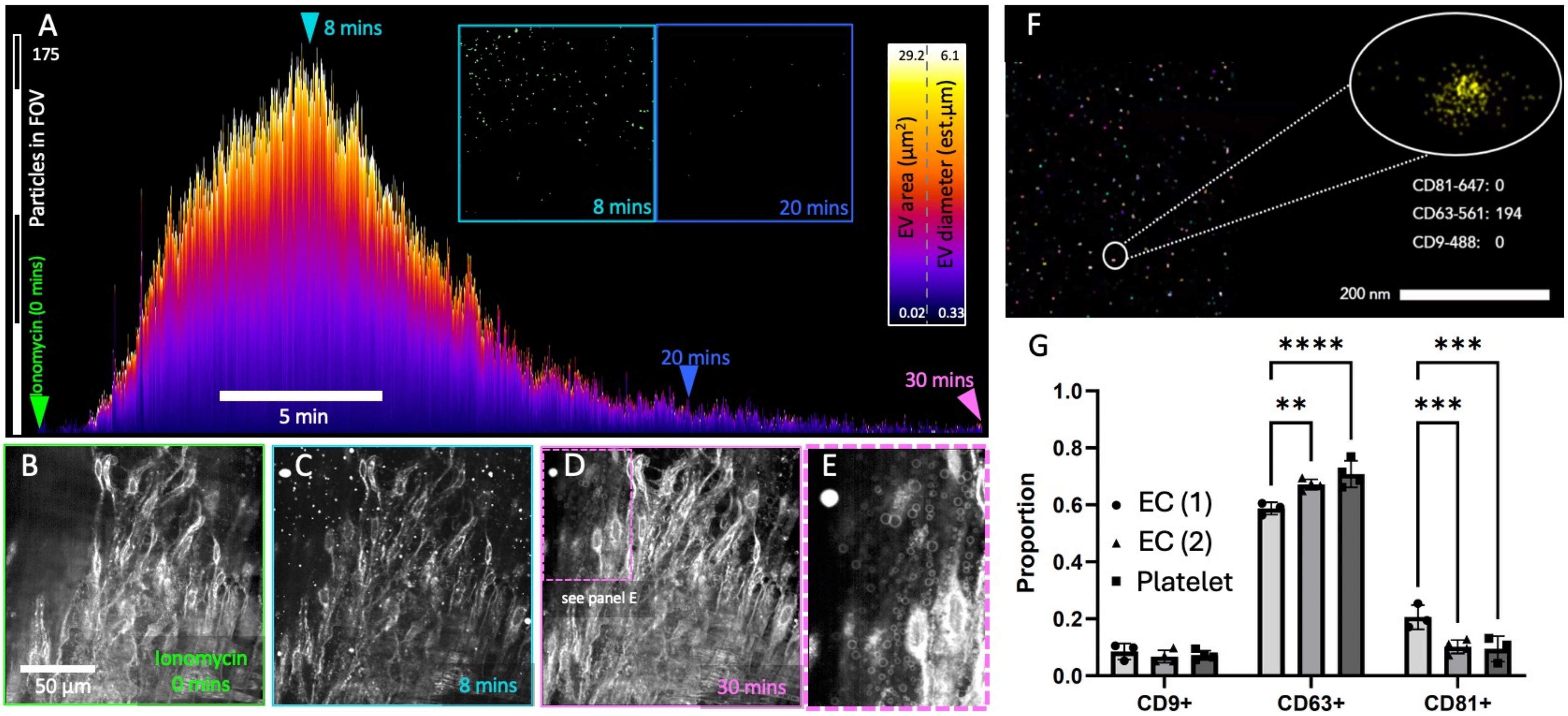
