## Supplemental figures for "Charge Screening by Extracellular Ca²⁺ Shields Endothelial Cells from Histone Injury and Permits Membrane Repair"

#### Contents

- Supplemental Methods
- References cited here appear in the main reference list
- Supplemental Figures 1–5 (legends below; figures supplied as separate image files)
- Supplemental Videos 1–12 (legends below; videos supplied as separate media files)

### Supplemental Methods

#### Human subjects

Healthy human subjects were enrolled for collection of citrated blood samples under a protocol approved by the Institutional Review Board. Fresh platelets were isolated by centrifugation at 200xg for 10 minutes without brake. Platelet rich plasma was removed, leaving erythrocytes and buffy coat. 1/10th volume of ACD-A was added to the platelet rich plasma and allowed to rest for 10 minutes. Platelet rich plasma was centrifuged at 1000xg for 10 minutes without brake. Platelet poor plasma was removed and banked at -80° C. The platelet pellet was resuspended in HEPES-buffered Tyrode's solution (137 mM NaCl, 2.7 mM KCl, 2 mM MgCl2, 0.42 mM $NaH_2PO_4$, 5 mM glucose, 10 mM HEPES, pH 7.4) supplemented with 1uM prostaglandin-E1. Platelets were counted on a hematology analyzer (Vetscan HM5, Zoetis, Parsippany, NJ), pelleted by centrifugation at 1000xG for 10 minutes without brake, and resuspended at a count of 5x10⁽5)/µL in HEPES- Tyrode's supplemented with 2 mM $CaCl_2$. Platelets were allowed to release their contents for 30 minutes on the benchtop. The releasate supernatant was isolated by centrifugation at 16000xG for 3 minutes, banked at -80° C, and thawed prior to use.

#### Animal husbandry

Adult male wild-type C57BL/6J mice (Jackson Laboratories) were kept on a 12 h light/dark cycle with ad libitum access to food and water and were housed in groups of five. All studies were conducted in accordance with the Guidelines for the Care and Use of Laboratory Animals (National Institute of Health) and approved by the Institutional Animal Care and Use Committee of the University of Vermont.

#### Imaging in blood vessels

En face murine vascular endothelial preparations were utilized for imaging as previously described^(7,16). Briefly, mice were euthanized using 5% Isoflurane, 95% $O_2$, and decapitated. The mesentery was immediately harvested and placed in 4°C HEPES-buffered physiological saline solution (HEPES-PSS: 10 mM HEPES, 134 mM NaCl, 6 mM KCl, 1 mM $MgCl_2$, 2 mM $CaCl_2$, 7 mM glucose, pH 7.4). Mesenteric arteries (100-150 µm internal diameter) were excised in sections of approximately 500 µm, cut longitudinally to expose the lumen, and pinned flat with the endothelial surface exposed en face for imaging. For $Ca^{2+}$ imaging experiments, vessels were loaded with green-fluorescent $Ca^{2+}$ indicator (5 µM Cal-520 AM, Thermo Fisher Scientific) and incubated at 37 °C for 30 minutes prior to imaging. For membrane imaging experiments the en face preparation was surgically injured using Vannas Spring Scissors (2 mm cutting edge; Fine Science Tools, Foster, CA) under 5.6X magnification (Olympus SZX7, Center Valley, PA). Vessels were then loaded with FM1-43 dye (2 µM, Thermo Fisher Scientific) at room temperature directly before imaging.

#### Scanning electron microscopy imaging of blood vessels

Third order mesenteric arteries (~200 µm diameter) from male C57BL6/j mice were cut open on one side longitudinally ("en face") and pinned down flat on a small silicone dish to expose the intact endothelium. Arteries were incubated with 50 µg/mL of unfractionated histones (Sigma-Aldrich, USA) for 30 minutes in HEPES-PSS (pH 7.4) at 37 °C. The arteries were then fixed in paraformaldehyde (2%)-glutaraldehyde (2.5%) cacodylate buffer solution (0.1 M; pH 7.2) for 24 h, washed with cacodylate buffer, post-fixed in a solution of 2.5% potassium ferrocyanide for 1 h, and washed again in cacodylate buffer and ultrapure water. Sections were dehydrated in ascending grades of ethanol, subjected to critical point drying in $CO_2$ (Samdri-PVT-3D, Tousimis, Rockville MD), coated with 10 nm of pure gold in a vacuum sputter coater (Quorum EMS150R S, Electron Microscopy Sciences, Hatfield, PA) and studied in a direct mode using a JEOL 6060 scanning electron microscope (Oxford Instruments, UK).

#### Cultured cells

Human endothelial (EA.hy926, ATCC) and kidney epithelial (HEK293, ATCC) cells were cultured according to manufacturer instructions. Genetically engineered HEK 293 cells with a triple knockout of ORAI1, ORAI2, and ORAI3 store-operated $Ca^{2+}$ entry (SOCE) channels generated using CRISPR-Cas9 gene editing as previously described28.

#### Live cell $Ca^{2+}$ imaging

$Ca^{2+}$ events were recorded in cells before and after histone stimulation using the $Ca^{2+}$ indicators. HEK cells were loaded with Fura-2 and imaged with a plate reader. Cells were manually identified and $Ca^{2+}$ levels within each cellular region of interest were reported as the ratio of $F_{340}/F_{380}$. ECs (EA.hy926 or mesenteric artery vascular preparations) were loaded with Cal-520 (5 µM) and imaged using spinning disk confocal microscopy. $Ca^{2+}$ oscillations from movies were analyzed using automated custom software (Volumetry, Grant Hennig) to create 2D and 3D prevalence maps allowing quantification of the numbers of cells activated per field and the frequency, amplitude, and duration of $Ca^{2+}$ events, as previously described16. Ionomycin was applied at the conclusion of each experiment as a positive control which maximally increased $Ca^{2+}$ levels in all cells within the field of view33.

#### Live cell membrane imaging

EA.hy926 cells were plated on glass coverslips and grown to ~70% confluence. The coverslips were transferred to individual wells containing HEPES-PSS with no added $Ca^{2+}$, 1.2 mM $Ca^{2+}$, or 12 mM $Ca^{2+}$ as indicated. Membrane dye FM1-43 (2 µM, Thermo Fisher Scientific) and $Gd^{3+}$ (10 µM) – where indicated – were added directly to the wells immediately prior to imaging. A baseline 3D Z-stack was then acquired before the careful addition of 1 mL extra HEPES-PSS (vehicle control), 50 µg/mL histones or 10 µM ionomycin. Continuous 2D recordings were acquired at 1 frame per second for 40 minutes. Finally, a second Z-stack was obtained. Individual cell cytoplasm ROIs were drawn manually in ImageJ, excluding outer membrane signal at all timepoints. Fluorescence intensity was quantified for each cell as ΔF/F0 using the following formula: (F_(t) - F_(0min)) / F_(0min). For representative Z-stack images, .png files were loaded into ImageJ and converted to 8-bit. Contrast was enhanced by lowering the maximum to 150 for each image within ImageJ. Field-of-view width was 490 µm, and magnified sub-panel width was 70 to 97 µm.

#### Live cell viability imaging

EA.hy926 cells were seeded at a density of 800,000 on 12 mm glass coverslips and allowed to grow for 1-2 days prior to imaging. Hoechst 33342 (1 µg/mL) and SYTOX Green (50 nM) were added to each well and incubated for 10 minutes at 37 °C in 5% $CO_2$. Coverslips were washed three times with HEPES-PSS following staining. For baseline and post-stim timepoints, three fields were imaged per coverslip, each consisting of a z-stack of 50 slices covering approximately 15-20 µm in depth. Following baseline imaging, coverslips were transferred into stimulation wells containing either HEPES-PSS control, histones (100 µg/mL) in HEPES-PSS, or ionomycin (10 µM) in HEPES-PSS. Cells were incubated at 37 °C for 1 hour. At 50 minutes, SYTOX Green was added to the stimulation wells and returned to the incubator. At the 1-hour mark, cells were imaged immediately (within 2-4 minutes), acquiring three non-overlapping fields per coverslip. After imaging, coverslips were washed and transferred into a well pre-filled with 2 mL of media and returned to the incubator for a 3-hour recovery period. After the 3-hour recovery in media, coverslips were removed, washed with HEPES-PSS, and placed in fresh HEPES-PSS containing SYTOX AADvanced for 10 minutes before final imaging. For the final timepoint, three fields per coverslip were imaged using a reduced stack depth of three slices (~15 µm). Cells were classified according to nuclear fluorescence at the final imaging time point, with SYTOX Green$^{+}$/SYTOX AADvanced$^{+}$ cells classified as permeable and without either marker classified as impermeable. SYTOX Green$^{+}$/SYTOX AADvanced$^{-}$ cells were classified as recovered, indicating permeability during the initial injury period followed by restoration of dye exclusion. SYTOX Green$^{-}$/SYTOX AADvanced$^{+}$ cells were not identified under any conditions. Percentages were calculated relative to the total number of Hoechst-positive nuclei detected at the final time point. Because different fields were sampled at each time point, cells lost through detachment during recovery were not included in the final analysis.

#### Recovery Imaging Analysis

Raw images were acquired using Fusion software and post-processed in Imaris. Standard parameters were applied uniformly across all replicates to ensure consistency, including four iterations of deconvolution and baseline subtraction using a 0.218 µm spatial threshold, as specified in the software's baseline correction settings. Nuclear regions were identified based on the Hoechst 33342 signal. Using the Imaris classification tool, each nucleus was then assigned to a viability category – double-positive (SYTOX Green$^{+}$/SYTOX AADvanced$^{+}$), SYTOX Green$^{+}$ only, SYTOX AADvanced$^{+}$ only, or negative – based on fluorescence intensity and channel overlap within the defined nuclear region. This approach ensured each cell was counted once and accurately categorized based on nuclear uptake alone. For display, red and green channel intensities were held constant across all images so that thresholding was consistent. The percentage of cells in each phenotype per coverslip was compared between recovery media by two-way ANOVA with Tukey correction for multiple comparisons.

##### Flow Cytometry

Cultured ECs were plated in 24-well tissue culture plates. Once cells were grown to confluency, the media was aspirated off and replaced with HEPES-PSS buffer solution containing histones as indicated. HEPES-PSS solutions contained no added $Ca^{2+}$, 1.2 mM $Ca^{2+}$, or 12 mM $Ca^{2+}$ depending on the condition. For membrane permeability assays, FM1-43 (2 µM) was added concurrently with histone exposure and incubated with the cells for 40 minutes at 37 °C with 5% $CO_2$. For experiments evaluating cell viability, cells were exposed to histones in HEPES-PSS containing 0, 1.2, or 12 mM $Ca^{2+}$, as indicated. Following treatment, cells were trypsinized, washed with PBS, and resuspended in $Ca^{2+}$-containing Annexin V binding buffer. All samples were stained with Annexin V (Biolegend #640920) and propidium iodide (Thermo Fisher #2301111) under identical conditions, independent of the extracellular $Ca^{2+}$ concentration during histone exposure. Cells were incubated at room temperature for 15 minutes before a 1:5 dilution in binding buffer. Samples were run on a flow cytometer (MACSQuant Analyzer 10, Miltenyi Biotec). Overall sample fluorescence was quantified as the median fluorescent intensity (MFI); n of 3-4 for all conditions.

##### Electric Cell-substrate Impedance Sensing

EA.hy926 cells were grown to confluence at a density of 200,000 cells on 8-well polyethylene terephthalate (PET) electrode arrays (8W10E+, Applied Biophysics, Inc., Troy, NY) for 48 hours.

Prior to seeding the electrode arrays were treated with 10 mM L-cysteine electrode-stabilizing solution for 15 minutes and allowed to air dry to provide a robust cleaning of the gold plating. Electric Cell-substrate Impedance Sensing (Model Z Theta, Applied BioPhysics, Inc., Troy, NY) data was collected from the capacitance values at 4,000 Hz over the course of 1.5 hours and normalized to the first measurement point.

#### Collection and Superresolution Microscopy of EVs

Extracellular vesicle-enriched preparations from human platelets or cultured endothelial cells (EA.hy926) were collected and characterized using super-resolution imaging. EA.hy926 cells were stimulated in serum free media with either HEPES-PSS, histones (50 μg/mL) or ionomycin (10 μM) for 1 hr at 37° C. Serum free media effluent was collected and filtered through 0.22 μm and 0.45 μm syringe tip filters and spun for 30 min at 4000xG at 4° C using Amicon Ultra 2 mL tubes (100,000 MWCO; Sigma-Aldrich, St. Louis, MO). 1 mL of EV-enriched preparation was collected and used for downstream analysis. EVs were labeled for canonical tetraspanin markers CD9, CD63, and CD81, as previously described.^(69,70) Briefly, a temperature-controlled Nanoimager S Mark II microscope (Oxford Nanoimaging) equipped with a 100X, 1.4NA oil immersion objective was used for direct stochastic optical reconstruction microscopy (dSTORM) of individual EVs labeled using the EV Profiler Kit (version 1; Oxford Nanoimaging), per manufacturer's instructions. CD9-AlexaFluor 488, CD63-AlexaFluor 568, and CD81-AlexaFluor647 antibodies from the kit served as fluorescent indicators for surface marker analysis.^(69,70)

#### Protein-Lipid Binding Assay

Mega Lipid Strips (Echelon-Inc, Salt Lake City, UT) kits were used in binding assays according to the manufacturer's instructions. Briefly, strips were blocked for one hour at room temperature with 3% bovine serum albumin (BSA: Sigma-Aldrich, St. Louis, MO) in PBS-T (phosphate buffered saline with 1% Tween-20). Strips were then incubated with Histone H3-GST (0.5 μg/mL; BPS Biosciences, San Diego, CA) for one hour at room temperature on an orbital shaker (170 RPM). Strips were then washed in PBS-T three times for 7 minutes each wash. Strips were then incubated in 75 μL of a GST secondary antibody (Echelon-Inc, Salt Lake City, UT) for one hour at room temperature. Strips were washed twice in PBS-T and twice in PBS for 5 minutes each. Strips were then incubated in 2 mL of TMB precipitating solution (Echelon-Inc, Salt Lake City, UT) for 2 minutes before imaging. Buffer containing $Ca^{2+}$ alone (nominal, 1.2 mM, or 12 mM) or $Ca^{2+}$ (1.2 mM) with $Gd^{3+}$ (30 μM) was added during the blocking step prior to the addition of histones where indicated. Following background subtraction, dot blot intensity was then quantified as area under the curve (AUC) using ImageJ software. The membrane lipid strips carried cardiolipin, PtdIns(4)P, phosphatidic acid, phosphatidylethanolamine, PtdIns(4,5)$P_2$, phosphatidylserine, phosphatidylcholine, sphingomyelin, and lysophosphocholine.

#### Data Analysis and Statistics

GraphPad Prism software (ver. 10.4.0; GraphPad Software, USA) was used for x–y graphing and analysis. Data were tested for normality using the Kolmogorov-Smirnov test. Parametric t-test or ANOVA, or appropriate non-parametric alternatives were applied for comparisons between groups, and differences were considered significant if $P < 0.05$.

#### $Ca^{2+}$ Fluorescence (Fura-2) Intensity

Max peak intensity was quantified for each cell for the period of stimulation. Since multiple measurements were taken from the same cell and compared across multiple stimuli, matched values for each cell were stacked. For each trace, a two-way ANOVA was conducted to compare the means across WT and ORAI TKO for each period of stimulation. Geisser-Greenhouse correction was used for non-sphericity and Šidák correction was used to correct for multiple comparisons.

#### Spatiotemporal Analysis of $Ca^{2+}$ Fluorescence (Cal-520)

Data was quantified as the total area with $Ca^{2+}$ signals, normalized as a percent of the ionomycin response. Thus, each independent replicate generated a percentage for baseline, $Gd^{3+}$, and histones. Two individual Friedman Tests for nonparametric, matched data were conducted for the cultured and en face preps respectively.

#### Lipid Binding

The area under the curve (AUC) was quantified in ImageJ for each lipid dot. A two-way ANOVA was used to compare histone-lipid binding for each lipid across different ionic conditions (0 mM $Ca^{2+}$, 12 mM $Ca^{2+}$, 30 μM $Gd^{3+}$). Data points were stacked by replicate (n=3). Geisser-Greenhouse correction was used for non-sphericity and Tukey correction was used to correct for multiple comparisons.

#### Membrane Imaging

Cell fluorescence (FM1-43) at 40 minutes was quantified as $\Delta F\_(40min)/F0$. All individual cell values were combined across replicates (n=3) and a Kruskal-Wallis test for non-parametric data was performed. A threshold of greater than 1.0 $\Delta F\_(40min)/F0$, representing 2-fold increase in fluorescence relative to baseline, was used to calculate the percent of cells filled with dye at 40 minutes.

#### Flow cytometry analysis

MFI was quantified using the median fluorescent value for each histone replicate (n=3 or 4) and compared across ionic conditions (0 mM, 1.2 mM, 12 mM $Ca^{2+}$) using a Kruskal-Wallis test.

#### ECIS analysis

Raw traces were normalized to the starting impedance at the time of histone treatment (time 0) so all traces aligned at 1.0. Endpoint resistance was taken at the very end of the experiment across all groups and compared using a t-Test. Total resistance area under the curve (AUC) for each run was automatically calculated using Graphpad Prism and compared using a t-test.

#### Supplemental Figure Legends

Supplemental Figure 1. Low-dose $Gd^{3+}$ does not block histone-induced $Ca^{2+}$ oscillations, and extracellular $Ca^{2+}$ is required. Wild-type (left) and ORAI triple-knockout (right) HEK-293 cells loaded with Fura-2 and stimulated with histones (50 µg/mL). Ionomycin (10 µM) applied at the end of each experiment as positive control. (A) $Gd^{3+}$ (5 µM) pre-treatment followed by histones, with quantification of maximum peak intensity per cell. (B) $Gd^{3+}$ (5 µM) post-treatment. (C) Histones applied in 0 mM $Ca^{2+}$ followed by restoration of 2 mM $Ca^{2+}$. Two-way ANOVA.

Supplemental Figure 2. Moderate-dose histones produce a subtle pre-apoptotic phenotype modulated by extracellular $Ca^{2+}$. EA.hy926 cells were treated with histones (50 µg/mL) in HEPES-PSS containing 0, 1.2, or 12 mM $Ca^{2+}$ for 1 h, then stained with Annexin V and propidium iodide. (A) Representative histograms of Annexin V (top) and propidium iodide (bottom) fluorescence for cells in buffer alone (blue) or with histones (pink), at each $Ca^{2+}$ concentration. (B, C) Median fluorescence intensity for Annexin V and propidium iodide. (D) Representative dot plots with the gating strategy. (E, F) Percent $PS^{+}/PI^{-}$ and $PS^{+}/PI^{+}$ cells. At this moderate dose the changes are smaller than those produced by 100 µg/mL histones (Figure 6), but follow the same dependence on extracellular $Ca^{2+}$.

Supplemental Figure 3. Low extracellular $Ca^{2+}$ converts the histone barrier response from transient stabilization to rapid failure. (A) Raw ECIS traces; histones (100 µg/mL) caused a rapid fall in monolayer resistance in low $Ca^{2+}$ but not in 1.2 mM $Ca^{2+}$. (B) Normalized endpoint resistance (1.7±0.1, n=8 versus 0.5±0.0, n=7). (C) Total resistance (AUC 1558±81, n=8 versus 462±9, n=7).

Supplemental Figure 4. Detailed experimental design for the membrane repair assay. (A) Schematic overview of the injury, staining, and recovery sequence. (B) Phenotype classification by dye combination.

Supplemental Figure 5. Extracellular vesicle release and reuptake in native tissue and vesicle surface marker characterization. The endothelial surface of a surgically opened vessel was mechanically injured with scissors and stained with FM1-43. Vesicle release was stimulated in remaining healthy cells with ionomycin (10 μM) and vesicles quantified by particle analysis of video data (A); the spatiotemporal histogram shows time on the x-axis (30 min) and vesicle count per field of view on the y-axis using a moving background average subtraction of ±10 s, with color indicating vesicle size as area ($\mu m^2$) or derived circular diameter (μm). (B–D) Time-lapse images of the field of view; injured cells appear bright, with uninjured cells visible at upper left. (E) The same cells demonstrate blebbing by the end of the recording. (F) Super-resolution (dSTORM) imaging of vesicles from stimulated ECs labeled for CD9, CD63, and CD81. (G) Summary of vesicle patterns produced by ECs exposed to ionomycin (1) or histones (2) compared with vesicles released by activated human platelets.

#### Supplemental Video Legends

Supplemental Video 1. Histones induce $Ca^{2+}$ influx in en face murine ECs. 3D representation of cumulative $Ca^{2+}$ oscillations over the 5 minute period after exposure to histones (50 μg/mL). Spatio-temporal representation overlaid on 2D image of the same field of view with ionomycin-induced maximal $Ca^{2+}$ response.

Supplemental Video 2. Pretreatment with $Gd^{3+}$ blocks histone-induced $Ca^{2+}$ influx in en face murine ECs. 3D representation of cumulative $Ca^{2+}$ oscillations after exposure to histones (50 μg/mL) in the presence of $Gd^{3+}$ (10 μM). Spatio-temporal representation overlaid on 2D image of the same field of view with ionomycin-induced maximal $Ca^{2+}$ response.

Supplemental Video 3. Video recording (25x) of ionomycin-induced endothelial cell membrane changes visualized using the membrane dye FM1-43.

Supplemental Video 4. Video recording (25x) of histone-induced endothelial cell membrane deformation visualized using the membrane dye FM1-43.

Supplemental Video 5. Full field of view with inset video of endothelial cell filopodia extrusion and movement after application of ionomycin (30 minute recording: speed 96x).

Supplemental Video 6. Full field of view with inset video of endothelial cell membrane blebbing after application of ionomycin (30 minute recording, speed 96x).

Supplemental Video 7. Full field of view with inset video of endothelial cell membrane ruffles after application of histones (20 minute recording, speed 96x).

Supplemental Video 8. Full field of view with inset video of endothelial cell rounding after application of histones (20 minute recording, speed 96x).

Supplemental Video 9. Full field of view with inset video of endothelial cell membrane blebbing after application of histones (14 minute recording, speed 96x).

Supplemental Video 10. Video of histones-induced endothelial cell membrane ruffling (30 minute recording, speed 96x).

Supplemental Video 11. Video of histone-induced endothelial cell membrane blebbing (30 minute recording, speed 96x).

Supplemental Video 12. Video of ionomycin-induced release and reuptake of extracellular vesicles (30 minute recording, speed 96x).
